# Oil biodegradation and marine oil snow formation in Arctic seawater microcosms at 4 °C

**DOI:** 10.64898/2026.11.02.763557

**Authors:** Thamali Vidanage, Durell Desmond, Dustin Isleifson, Gary Stern, R. Eric Collins

## Abstract

Hudson Bay is a sub-Arctic inland sea characterized by seasonal sea-ice cover, which governs marine circulation, biological productivity, and ecosystem structure. In recent decades, the duration of sea-ice cover has declined, resulting in longer ice-free seasons that are expected to elevate marine activity and environmental exposure to anthropogenic disturbances. As such, the potential for oil spill incidents associated with increased marine operations is expected to rise. To understand microbial biodegradation dynamics associated with marine oil snow (MOS), seawater microcosms amended with diesel fuel were incubated at 4 °C for 6 weeks. We amended all our microcosms with UV-weathered Churchill diesel fuel at 1% (v/v). 16S rRNA sequencing of the incubations revealed a decline in overall microbial diversity and a dramatic increase in specialized oil-degrading taxa, with *Pseudomonas* and *Pseudoalteromonas* being the main taxa involved, as both effectively degrade hydrocarbons under ambient and stimulated conditions. Biostimulation with nutrients (NaNO_3_, KH_2_PO_4_) and an alternative dispersant at 0.1% (v/v) resulted in a greater abundance of MOS aggregates and rapid enrichment of oil-degraders. Simple n-alkanes exhibited the highest biodegradation loss relative to complex polycyclic aromatic hydrocarbons (PAHs); however, biostimulation did not substantially enhance their degradation. Additionally, UV weathering resulted in a 22% loss of PAHs, which is three times higher than the UV loss observed for n-alkanes. Biostimulated samples exhibited an elevated transfer of high molecular weight alkanes and PAHs from the oil into the aqueous phase, accompanied by a rapid enrichment of oil-degrading taxa.

## 1 Introduction

The Arctic is becoming more accessible as a result of the recent decreases in sea ice coverage brought about by climate change (Scheibye et al., 2017). In the last five decades, Arctic sea ice extent has decreased by 5 to 20% each decade, particularly during the summer months (Mudryk et al., 2021). The Canadian Arctic is a pivotal region for trans-Arctic shipping, with interest gaining in transit through the Northwest Passage (Mudryk et al., 2021). Between the years 2013 and 2023, the distance traveled by ships in the Arctic has doubled (PAME, 2024), with fishing and cargo shipments being the major drivers of the increased vessel traffic (Eguíluz et al., 2016; Pizzolato et al., 2016).

Hudson Bay experiences longer ice-free seasons than the high Arctic (Stroeve et al., 2024), facilitating increased shipping access and traffic in the region (Andrews et al., 2017). Sea ice extent in Hudson Bay in the spring of 2024 reached a new record low for the month, falling below the median from 1979 to 2023 (Soriot et al., 2025), raising alarms regarding the potential oil spill risks associated with shipping. The Port of Churchill is the only deep-water port in the Canadian Arctic, and it provides access to North America via the Hudson Bay railway (Andrews et al., 2018). Despite its closure in 2016, the 2025 federal budget shed light on developing transformative energy and trade corridors by expanding the export capacity of the Port of Churchill to strengthen trade with Europe and other partners, and improve connectivity between Churchill, Manitoba, and the rest of Canada (Department of Finance Canada, 2025). Shipping in Hudson Bay and Hudson Strait generated by the mining industry is expected to grow quite rapidly in the coming years (Andrews et al., 2017), while the majority of the shipping traffic in Hudson Bay appears to be concentrated along the corridor across the Bay to Churchill (Judson, 2010). Though shipping can improve the livelihoods of communities by delivering essential supplies, it also carries risks that can affect the marine environment, including oil spills, the introduction of invasive species, and disturbances to the migration routes of Arctic mammals. Ultimately, these economic activities most affect the Inuit communities in the regions that rely on harvesting country foods from the natural environment for subsistence (Olsen et al., 2019; Dawson et al., 2020; Lasserre, 2022; Adeniran-Obey and Osagie, 2024).

In aerobic marine waters, the majority of hydrocarbons are degraded by microbes that use oil as their source of energy and carbon, with a half-life of days to months (Hazen et al., 2016). Nonetheless, rates of biodegradation depend on the oil composition, state, and concentration (Leahy and Colwell, 1990). Even though psychrophilic (cold-loving) oil degraders adapt to low temperatures with specialized enzymatic pathways (Ribicic et al., 2018), removal of oil from biodegradation alone is challenging due to freezing temperatures that slow down the recovery (Garneau et al., 2016; Stern et al., 2023), thus the impact on marine life, communities, and socioeconomic conditions can be long-lasting (Bi et al., 2025).

Dispersed oil undergoes a range of physical, chemical, and biological processes, such as evaporation, dissolution, photochemical oxidation, biodegradation, and sorption onto particulate matter in seawater (Zhao et al., 2016). Under certain conditions, interactions between dispersed oil, organic matter, minerals, and microorganisms promote the formation of marine oil snow (MOS) that eventually sinks to the sea floor (Gregson et al., 2021; Ross et al., 2021). MOS may develop as pre-formed marine snow (MS) interacting with oil, or oil acting as an agent for microbial biofilm growth and floc formation (Gregson et al., 2021), and host a distinct bacterial community different from the free-living community in the surrounding seawater (Arnosti et al., 2016). These aggregates transfer oil, and extracellular polymeric substances (EPS)-derived dissolved organic carbon (DOC) into the water column and contribute to the carbon flux through oil degradation (Ziervogel et al., 2012). The formation and aging of MOS foster the establishment of specialized microbial communities that enhance the degradation of oil and oil-derived residues (Arnosti et al., 2016). As aggregates descend, microbial degradation preferentially removes labile hydrocarbons, enriching the residue in less bioavailable compounds (Gregson et al., 2021).

Sinking MOS aggregates deliver oil and oil residues to deeper marine environments, where distinct microbial communities further transform less bioavailable hydrocarbons (Quigg et al., 2020). MOS can potentially impact the benthic organisms through direct toxicity and reduced oxygen availability due to microbial respiration (Yang et al., 2016; van Eenennaam et al., 2018). Therefore, understanding the rate at which hydrocarbon composition changes during transit through the water column and subsequent sedimentation is critical for evaluating potential risks to benthic communities (Atlas and Hazen, 2011).

The Arctic region is expected to be particularly sensitive to MOS formation due to unique environmental conditions, including low temperatures and the presence of sea ice, which together result in slower biodegradation rates (Murk et al., 2020). Diesel fuel is a complex mixture of hydrocarbons that includes both aliphatic compounds, such as alkanes, and aromatic compounds, including polycyclic aromatic hydrocarbons (PAHs; Liang et al., 2005). Aliphatic hydrocarbons constitute roughly 64% of diesel fuel, with approximately 35% including both monoaromatic compounds and PAHs. Olefinic hydrocarbons typically represent a minor fraction, comprising approximately 1–2% of the total composition (Gad, 2014).

Both diesel and crude oil contain hundreds of PAHs plus alkylated PAH compounds that can be more persistent than their parent compounds (Latimer and Zheng, 2003) and exhibit varying degrees of hydrophobicity depending on the number of aromatic rings (Sakellari et al., 2021). The United States Environmental Protection Agency (USEPA) has listed 16 PAHs as priority pollutants given their potential toxicity, including naphthalene, acenaphthylene, acenaphthene, fluorene, phenanthrene, anthracene, fluoranthene, pyrene, benz[a]anthracene, chrysene, benzo[b]fluoranthene, benzo[k]fluoranthene, benzo[a]pyrene, indeno[1,2,3-cd]pyrene, dibenzo[a,h]anthracene, and benzo[ghi]perylene (USEPA, 2014). Generally, PAHs have greater resistance to degradation than other compound groups such as alkanes (Friedman et al., 2014). PAHs associated with oil spills are expected to be fully bioavailable to marine fauna (Yunker et al., 2015), with some being carcinogenic and mutagenic (Latimer and Zheng, 2003).

Despite extensive research on hydrocarbon biodegradation, uncertainties remain regarding how nutrient enrichment and dispersant applications influence the fate of different hydrocarbon classes and associated microbial communities. In particular, the combined effects of biostimulation on the degradation of n-alkanes and PAHs, as well as on the selective enrichment of hydrocarbon-degrading microorganisms, are not fully understood under environmentally relevant conditions. In addition, oil–microbe interactions can promote the formation of MOS, which plays a critical role in the aggregation, vertical transport, and microbial processing of oil in the water column. However, the extent to which nutrient and dispersant additions modulate MOS formation and its associated microbial consortia, and how these processes influence hydrocarbon biodegradation and partitioning remain poorly understood.

In previous work, we characterized short-term microbial responses to differential oil exposure and biostimulation within artificial seawater microcosms over one week, capturing early community shifts but not longer-term succession or sustained hydrocarbon transformation processes. In contrast, the present manuscript extends this work by examining microbial community dynamics and biodegradation patterns over six weeks under more Arctic-relevant conditions. This longer experimental duration enables evaluation of temporal restructuring, treatment persistence, and the sustained enrichment of hydrocarbon-degrading taxa, providing insight into longer-term ecosystem responses to oil contamination. This study aimed (1) to elucidate the processes underlying MOS formation in the Canadian Arctic, (2) to characterize microbial community shifts in a low-temperature seawater microcosm, and (3) to evaluate hydrocarbon compositional changes and phase partitioning under biostimulation.

## 2 Materials and methods

### 2.1 Field sampling

Seawater samples were collected using a peristaltic pump into a large sterile cubitainer from the Control pool at the Ocean Sea Ice Mesocosm (OSIM) facility of the Churchill Marine Observatory (CMO) on 21^st^ April 2025 (Figure 1). The water had been previously pumped into the mesocosm pools from the Churchill estuary on October 16, 2024; the salinity was 26.4 PSU.

**Figure 1.**
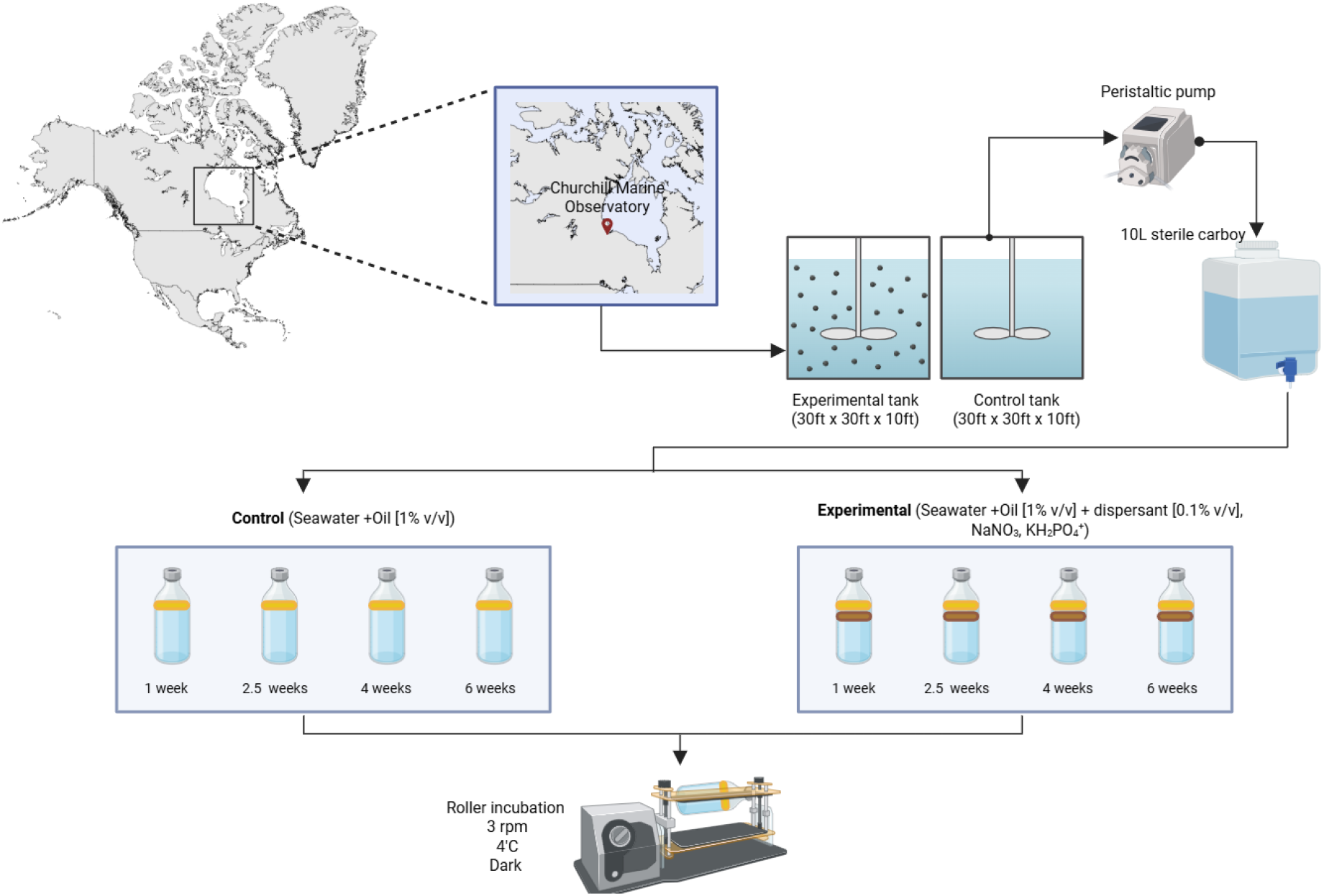
Water sample collection from CMO and roller incubation setup at 3 rpm rotation speed and at 4 °C in the dark. Created with BioRender.com.

Upon return to the University of Manitoba, the water sample was stored at 4 °C in the cold room facility at the Centre for Earth Observation Science for 2 weeks until the microcosms were set up.

### 2.2 Microcosm setup and subsampling

The microcosms were established in 125 mL Pyrex glass bottles on three tube rollers, as previously described by Gutierrez et al. (2013). Two types of treatment groups were established, and each treatment was run in duplicate for a period of six weeks (Figure 1). Aliquots of Churchill diesel fuel were UV-weathered using UV-A (365 nm) and UV-B (302 nm) lamps for 5 days before the experiment. A total of 16 microcosms were set up, along with six additional samples for chemical analysis. Eight microcosms consisted of 98.9 mL of seawater amended with UV-weathered Churchill diesel fuel at a ratio of 1% (v/v), 0.1% (v/v) alternative dispersants (soy lecithin and Tween 80 dissolved in ethanol (Athas et al., 2014) at a 60/40 [w/w] ratio), and nutrients (7.785 mM NaNO_3_ and 0.2 mM KH_2_PO_4_) to satisfy the Redfield ratios. The other eight microcosms consisted of 99 mL of seawater and 1% (v/v) UV-weathered Churchill diesel fuel. All the microcosms were placed on the tube rollers horizontally, and the speed was set at 3 rpm at 4 °C to mimic the natural environmental conditions of the ocean. The treatments and controls were sampled at four time points over 6 weeks for microbial analysis: T0 at Day 0, T1 at 1 week, T2 at 2.5 weeks, T3 at 4 weeks, and T5 at 6 weeks. Observations were recorded at each sampling point.

### 2.3 Genomic DNA extraction

MOS that could be handled without breaking were grown on Bushnell-Haas agar medium (30% w/v NaCl) at 4 °C with motor oil as the sole carbon source. The majority of the MOS aggregates were too fragile to withdraw reliably (Figure 2). Hence, the whole sample was filtered onto a 0.22 *µ*m Sterivex filter for genomic DNA extraction. The Qiagen DNeasy PowerSoil kit was used to extract DNA from the samples following the manufacturer’s instructions, with slight modifications. For all extractions, the bead-beating tubes containing the samples and CD1 solution were heated to 70 °C for 10 minutes to facilitate cell lysis (Murphy et al., 2021). Additionally, an aliquot of elution buffer was heated to 50 °C before eluting the DNA in the final step. All extracted DNA products were quantified using a Qubit fluorometer and stored at −70 °C until the sequencing was performed.

**Figure 2.**
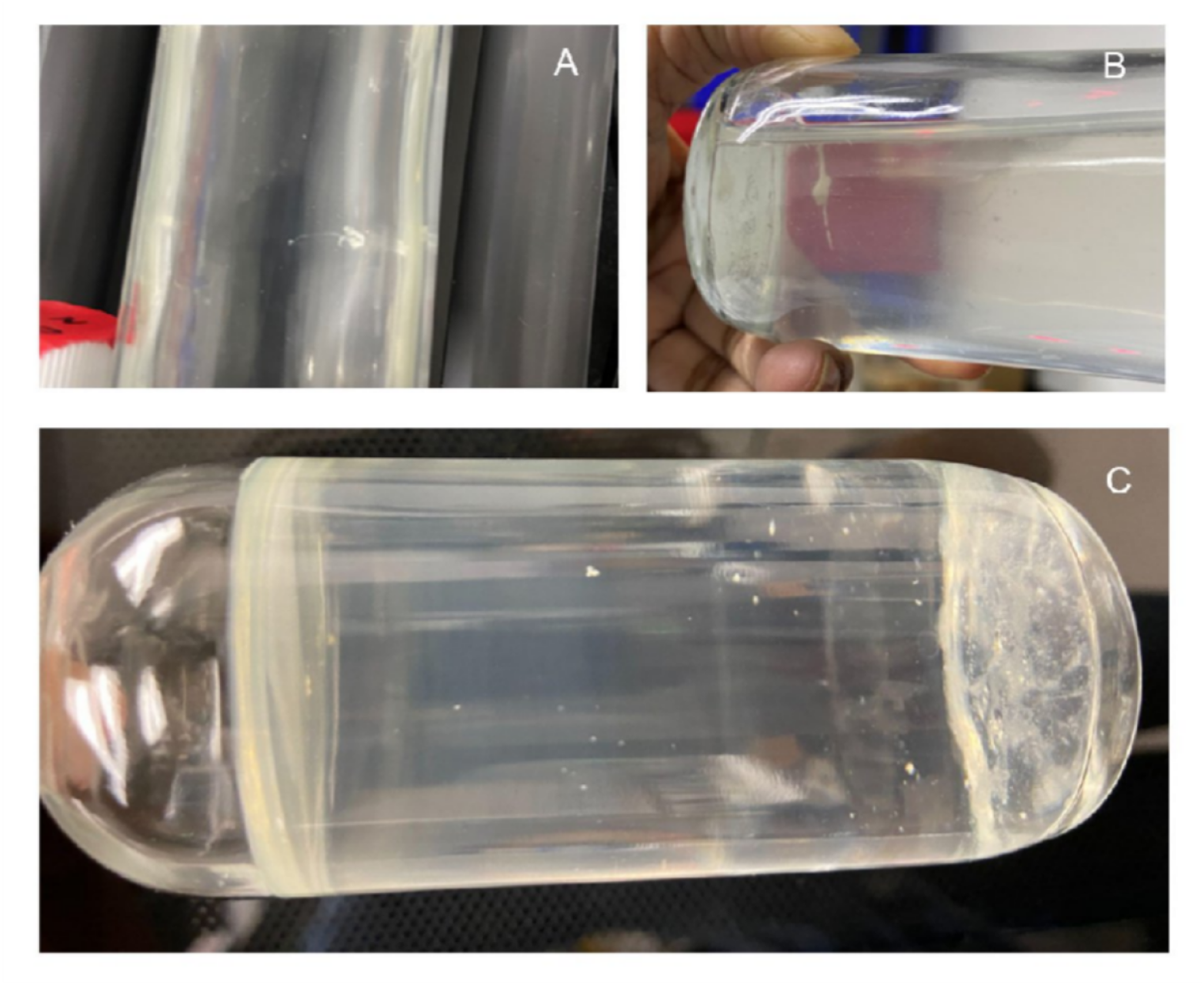
MOS formation in the biostimulated treatments. (A) A MOS formed in a biostimulated treatment group within a day. (B) The same MOS after 1 week of roller incubation. (C) The mucous MOS settled at the bottom of the bottle.

### 2.4 16S rRNA amplicon sequencing

Paired-end (2 × 250) amplicon sequencing across the V4 hypervariable region of the 16S rRNA gene was carried out on the Illumina NovaSeq platform using the 515F (GTGYCAGCMGCCGCGGTAA) and 806RB (GGACTACNVGGGTWTCTAAT) primer pair (CD Genomics, New York, USA). Approximately 605k high-quality reads were obtained with a read depth of 19,197 to 58,613 per sample. Raw sequencing reads were quality-controlled by removing adapters and primer sequences using Cutadapt v5.2 (Martin, 2011). Trimmed reads were then processed using the DADA2 (Callahan et al., 2016) package in R to infer amplicon sequence variants (ASVs), and taxonomic assignment was performed using the SILVA v138.2 reference database (Quast et al., 2013). Code for the bioinformatics pipeline is available at https://github.com/rec3141/microscape.

### 2.5 Hydrocarbon analysis by gas chromatography–mass spectrometry (GC-MS)

Hydrocarbons were extracted from the microcosms after 6 weeks. To account for the hydrocarbon fractions that were transported from the oil phase to the aqueous phase, each microcosm group was analyzed in three sub-sections: oil phase (consisting of two replicates), aqueous phase (consisting of two replicates), and whole sample (with no replicates) for both control and biostimulated treatments. The oil phase was decanted and measured volumetrically, and a homogenized aliquot (20 *µ*L) was diluted to 2%, 1%, and 0.1% with isooctane before chemical analysis. For the aqueous phase, each microcosm was extracted three times with hexane and dichloromethane (DCM) as solvents. Extracts were filtered through sodium sulphate to remove any residual water. The extracts were then evaporated on a BUCHI Rotary Evaporator with a 30 °C water bath, a starting pressure of 450 mbar, and a final reduced pressure of 150 mbar, until the extracts reached approximately 1 mL (in isooctane). For the whole sample, extraction and reduction proceeded identically to the aqueous phase but were instead reduced to a 10 mL hexane stock before further diluting by 5% and 3% with isooctane. For each final extract, a 2-fluorobiphenyl (2-FB) internal standard was added and the final sample volume was adjusted to 1 mL (by mass) in a GC vial. All chemicals used were Fisher Chemical Optima grade.

An Agilent 7010B Triple Quadrupole GC-MS (QQQ-MS) was employed for the analysis of aromatics and alkanes using multiple reaction monitoring (MRM). The system was equipped with an Rxi-PAH column (60 m × 250 *µ*m × 0.1 *µ*m). For aromatics, the oven temperature program was held at 40 °C for 1.5 minutes, then ramped to 140 °C at 30 °C/min. It was subsequently increased to 250 °C at 3 °C/min, followed by a third ramp to 320 °C at 2 °C/min, where it was held for 20 minutes. For alkanes, the oven temperature program was held at 30 °C for 2 minutes, then ramped to 120 °C at 15 °C/min. It was subsequently increased to 250 °C at 3 °C/min, followed by a third ramp to 320 °C at 2 °C/min, where it was held for 30 minutes. An electron ionization of 70 eV was used. The injection volume of each sample was 1 *µ*L, which was carried by helium gas at a flow rate of 1.1 mL/min through the GC column. Nitrogen was used as the collision gas at a flow rate of 1.5 mL/min and an ion gain of 5. 2-FB was measured in each sample to correct for instrument response.

Quantification of aromatics was carried out using a PAH calibration mixture composed of the following standards: Polycyclic Aromatic Hydrocarbons Mix (CV Calibration Mix #5, Restek) and PAH/Dibenzothiophenes Mixture (Chiron). All aromatic calibration curves consisted of at least 9 points and had R^2^ values ≥ 0.95. Quantification of alkanes was carried out using an n-alkane calibration mix consisting of C_8_–C_40_ n-alkanes (Sigma-Aldrich). All alkane calibration curves consisted of at least 8 points and had R^2^ values ≥ 0.98. A minimum signal-to-noise ratio of 5 was achieved for all compounds. A full list of the compounds analyzed, and their MRM quantifier settings, is included in the supplementary material (Table S1).

### 2.6 Diagnostic ratios for determining biodegradation

The extent of n-alkane and PAH biodegradation was determined from the abundance of heptadecane relative to pristane (n-C_17_/pristane) (Johnston et al., 2007) and 2-methylnaphthalene relative to 1-methylnaphthalene (2MN/1MN) (Murphy et al., 2021) in fresh diesel, UV-weathered diesel, biostimulated, and oil-only control treatments.

### 2.7 Statistical analysis

Microbial community analysis was performed using R. All plots were generated in R (v.4.3.1) using the ggplot2 package (Wickham, 2016). Statistical significance across the treatment groups was analyzed using two-tailed t-tests (*α* = 0.05).

## 3 Results

### 3.1 MOS development

The earliest time for MOS development in the roller incubations was less than 24 hours. Initially, MOS particles were suspended in the water column and were off-white in color and approximately 2 to 3 mm in size (Figure 2). However, over time, they aggregated and sank; the majority of MOS particles seen after 2.5 weeks were settled at the bottom, and only a very few tiny particles could be seen scattered in the water column. The MOS settled at the bottom had a mucus texture rather than being dense. The shapes varied from linear to round. Notably, MOS were observed only in the biostimulated (nutrient and dispersant added) treatments. No control treatment had MOS or even visible signs of MOS formation during the 6 weeks. Instead, the control treatment had the oil layer visible on the top of the water column when the bottle was inverted. Since the MOS particles were extremely fragile, they disintegrated quickly when inverting the bottles for sampling or photography.

### 3.2 Microbial community composition between treatment groups

Following quality filtering and denoising, ASVs were inferred, resulting in 243 ASVs spanning 10 phyla, 13 classes, 44 orders, 74 families, and 100 genera. A pronounced dominance of *Pseudoalteromonas* and *Pseudomonas* was observed at 2.5 weeks in the biostimulated treatments, with *Pseudoalteromonas* accounting for 82.5% of the total microbial abundance (Figure 3). However, the relative abundance of *Pseudoalteromonas* declined markedly over time, decreasing from 46.9% at four weeks to 11.2% at six weeks. In contrast, *Pseudomonas* exhibited a strong increase, rising from 52.2% at four weeks to 85.4% at six weeks.

**Figure 3.**
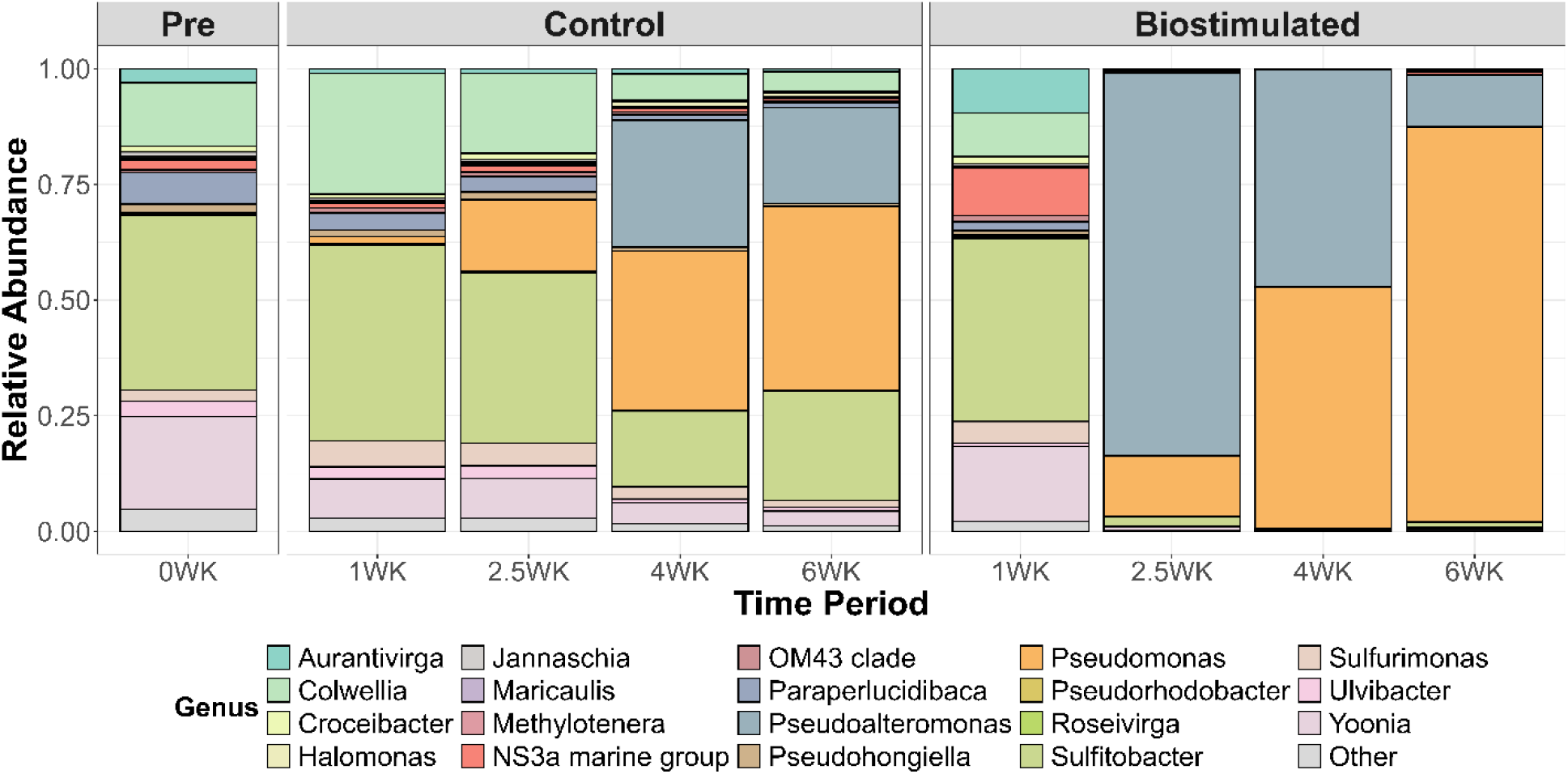
The mean genus-level composition of the microbial taxa present in the treatment groups (*n* = 2 per treatment). The taxa with relative abundance less than 0.001 were grouped as “other”.

The addition of nutrients and dispersants rapidly altered microbial community composition, compared to oil-only controls, favoring the proliferation of a limited number of oil-degrading taxa under these enriched conditions. In the control treatments, the relative abundances of *Pseudomonas* and *Pseudoalteromonas* also increased over time; however, a more diverse assemblage of oil-degrading taxa was observed compared to the biostimulated treatments. *Colwellia* was present in the initial seawater at 13.6% and increased to 26.1% after one week before subsequently declining. During the first 2.5 weeks of control treatments, *Sulfitobacter* was the dominant taxon, accounting for 37.7% of the community. As *Pseudomonas* populations increased, the relative abundance of *Sulfitobacter* declined. By four weeks, both *Pseudoalteromonas* and *Pseudomonas* had emerged as dominant taxa in control treatments. Nevertheless, *Pseudomonas* appeared to outcompete *Pseudoalteromonas*, ultimately becoming the predominant taxon after six weeks.

### 3.3 Biodegradation of hydrocarbon compounds

Compared to PAHs, n-alkanes showed a more pronounced loss in both control and biostimulated treatments. Nonetheless, biostimulation had a similar impact on the overall biodegradation of both n-alkanes (C_10_–C_23_) and PAHs as revealed by GC-MS (Figure 4). This is evident from the nearly similar ratio of n-C_17_/pristane (0.83) and 2MN/1MN (1.39) in both treatments (Figure S1). Notably, the above ratios were lower than those of the n-C_17_/pristane and 2MN/1MN ratios for fresh diesel (1.36 and 1.54) and UV-weathered diesel (1.42 and 1.53), suggesting active biodegradation processes in both treatments. Reduced n-C_17_/pristane and 2MN/1MN diagnostic ratios in control and biostimulated treatments relative to fresh and UV-weathered diesel, along with similar ratios between fresh and UV-weathered diesel, indicate comparable biodegradation patterns, as UV exposure primarily drives physicochemical weathering rather than biodegradation. A significant loss of PAHs was observed during the UV weathering process compared to the n-alkanes (*p* = 0.028; Figure S2). PAHs displayed compound-specific responses to UV-driven weathering, with the highest loss (68.2%) observed for pyrene (C_16_H_10_). Despite this, UV exposure did not produce a systematic or consistent alteration pattern among PAHs.

**Figure 4.**
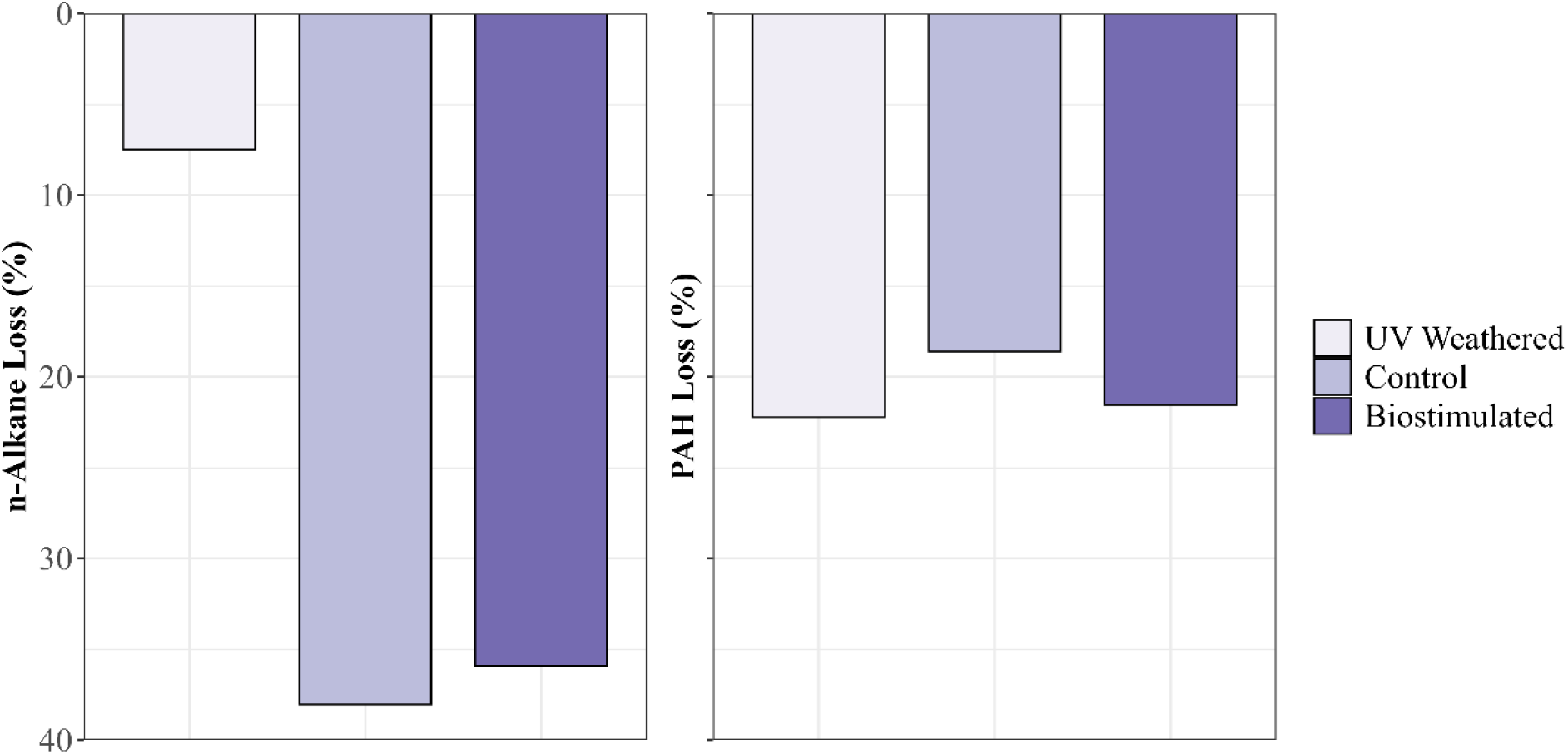
Total loss (%) of n-alkanes (a) and PAHs (b) due to UV weathering and biodegradation after 6 weeks of incubation.

### 3.4 Partitioning of hydrocarbon compounds

Higher oil-to-aqueous ratios for alkanes in the control treatments reflect the absence of enhanced partitioning processes (e.g., dispersant-mediated dispersion or incorporation into marine oil snow), resulting in a greater proportion of oil remaining in the bulk phase (Figure 5 and Figure S3). Within the controls, low-molecular-weight (LMW) alkanes exhibited higher ratios compared to high-molecular-weight (HMW) alkanes, likely due to the greater susceptibility of LMW compounds to biodegradation in the aqueous phase. A similar trend was observed for the biostimulated treatments in which LMW alkanes exhibited higher ratios compared to HMW, albeit less pronounced. These patterns are likely driven by enhanced partitioning of alkanes into the aqueous phase in the presence of dispersants and MOS aggregates. Among the partitioned n-alkanes, LMW compounds were preferentially biodegraded, whereas HMW n-alkanes exhibited higher retention in the aqueous phase through partitioning relative to their biodegradation rates.

**Figure 5.**
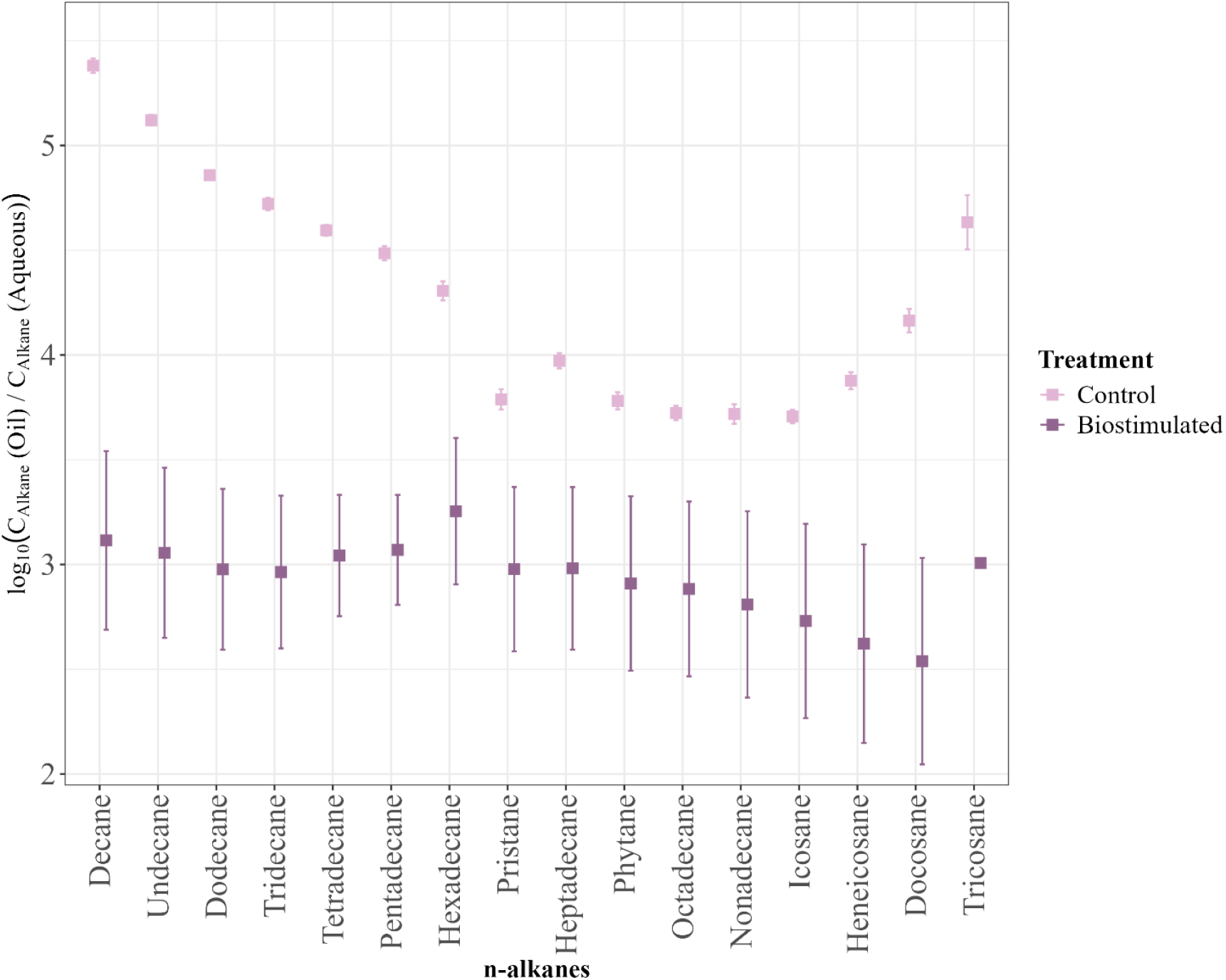
Partitioning of individual n-alkanes from diesel fuel between the oil and aqueous phases in control and biostimulated treatments after six weeks of incubation. Compounds are arranged in order of increasing molecular weight. The ratio of n-alkane concentrations in oil versus aqueous phases is shown on a log_10_ scale. Error bars represent standard deviations across replicates, and concentrations were normalized to total sample volume for direct comparison of partitioning behavior across treatments.

Consistent with these patterns, PAHs in the control treatments exhibited higher oil-to-aqueous ratios compared to the biostimulated treatments. Within the controls, HMW PAHs displayed comparatively lower ratios. Biostimulated treatments showed markedly reduced ratios relative to controls, reflecting enhanced biodegradation facilitated by increased partitioning of PAHs into the aqueous phase through MOS sorption and dispersant action, in which HMW PAHs in biostimulated treatments were retained in the water column to a greater extent than they were biodegraded, as indicated by the declining oil-to-aqueous ratios (Figure 6 and Figure S4).

**Figure 6.**
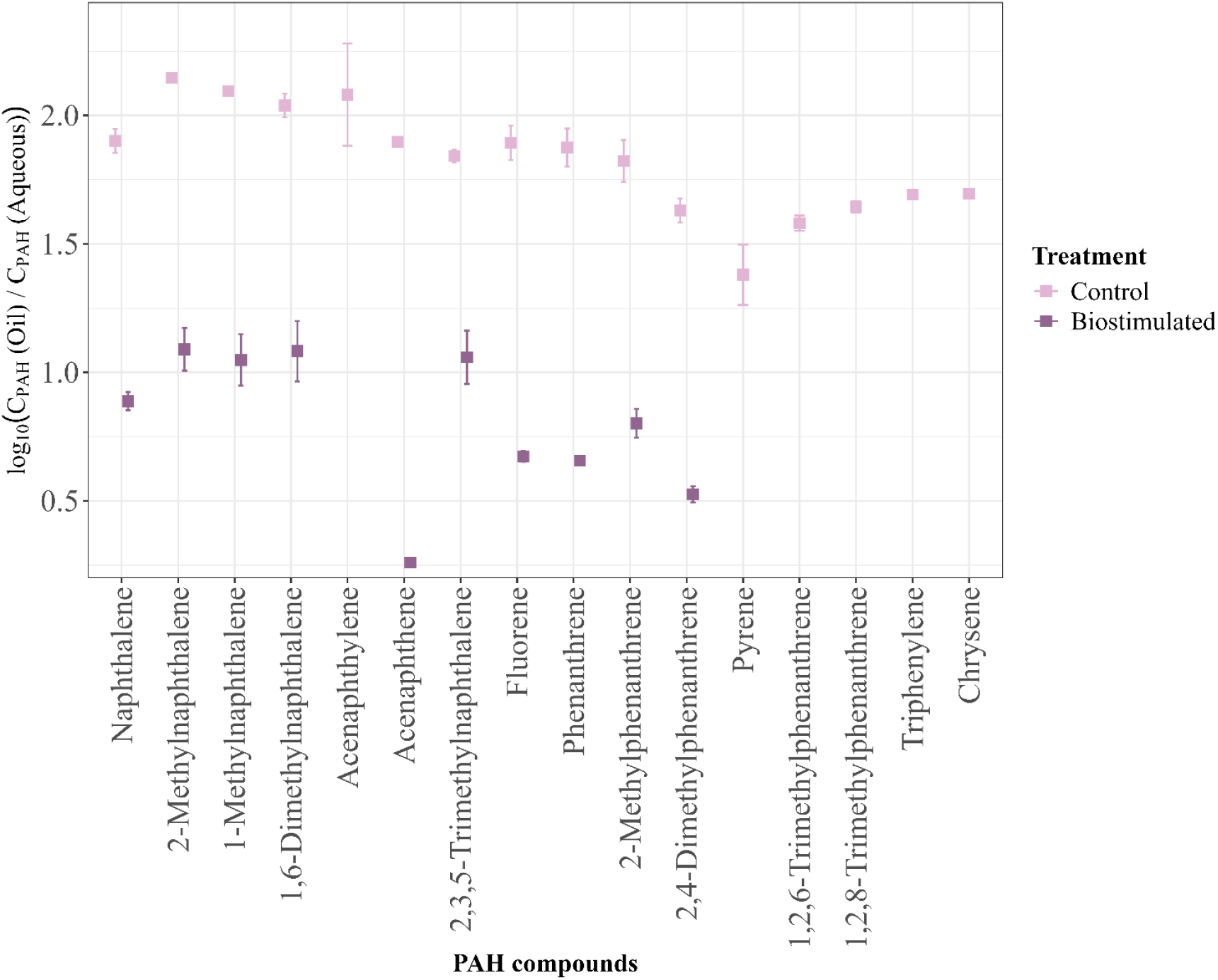
Partitioning of individual PAHs from diesel fuel between the oil and aqueous phases in control and biostimulated treatments after six weeks of incubation. Compounds are arranged in order of increasing molecular weight. The ratio of PAH concentrations in oil versus aqueous phases is shown on a log_10_ scale. Error bars represent standard deviations across replicates, and concentrations were normalized to total sample volume for direct comparison of partitioning behavior across treatments.

## 4 Discussion

### 4.1 Biostimulation influences the early succession of oil-degrading taxa

To our knowledge, this is the first study conducted on MOS formation from Canadian Arctic waters. Our roller bottle incubations indicate that MOS formation is independent of climate, as it has also been observed in other studies conducted in temperate and the Northeast Atlantic oceans (Passow et al., 2012; Fu et al., 2014; Passow, 2016; Suja et al., 2017). MOS formation was observed exclusively in biostimulated treatments containing nutrients and dispersants, where no aggregation occurred in oil-in-seawater controls. This suggests that nutrient enrichment and dispersant application facilitate oil-particle interactions and EPS production, which are central to MOS formation. Similar mechanisms have been reported following the Deepwater Horizon (DWH) spill, where microbial EPS production promoted rapid formation of oil-rich marine snow (Passow et al., 2012). This confirms that MOS formation is not an inherent property of oil alone, but a biologically mediated process sensitive to spill response strategies. Microorganisms play a vital role in the natural attenuation of oil spills, and bioremediation serves as a cost-effective approach in a wide range of ecosystems, including the Arctic (Aislabie et al., 2006; Góngora et al., 2022). In natural marine environments, biodegradation of petroleum hydrocarbons involves a succession of microbial consortia (Peeb et al., 2022). Following an oil spill, groups of hydrocarbon degraders, typically found in low relative abundance (<1%) in seawater, increase their numbers and become dominant as a response to the spillage (Neethu et al., 2019). We observed that oil-degrading bacteria proliferated even in control treatments upon the addition of oil, although the onset of colonization was delayed. Also, MOS formation was accompanied by shifts in microbial community composition relative to controls. These changes suggest the selective enrichment of hydrocarbon-associated and EPS-producing bacteria within MOS aggregates. We posit that the dramatic increase in *Pseudoalteromonas* population within 2.5 weeks of incubation was due to EPS production, which concurrently caused MOS formation. Similar observations have been reported from the oil-in-seawater microcosms conducted using North Atlantic seawater, where an increase in the abundance of *Pseudoalteromonas* in MOS particles was observed after 2.5 weeks (Suja et al., 2017). Species of *Pseudoalteromonas* are members of the “generalist” hydrocarbon degraders in the marine environments (Gutierrez et al., 2013) with the ability to produce EPS (Suja et al., 2017). These bacteria significantly contribute to the ocean’s pool of dissolved organic matter (DOM) during spill events. They enhance oil aggregation, dissolution, bioavailability, and the eventual degradation of hydrocarbons by indigenous bacterial communities (Gutierrez et al., 2013). In general, EPS provides a matrix for MOS formation (Passow, 2016), and nutrient-enriched treatments yield higher concentrations of EPS, highlighting the supportive role of nutrient enrichment for EPS-producing bacteria (Suja et al., 2017).

*Colwellia* populations did not increase in abundance in biostimulated treatments, despite being relatively abundant in control treatments within 2.5 weeks. One possible explanation is *Colwellia*’s preference for degrading gaseous and simple hydrocarbon aromatics (Mason et al., 2014) that are commonly present in fresh petroleum products (Reddy et al., 2012). As oil undergoes weathering, these volatile and gaseous components are progressively lost through evaporation, dissolution, and biodegradation (Tarr et al., 2016). The depletion of these preferred substrates provides a plausible explanation for the decline in *Colwellia* relative abundance observed after approximately 2.5 weeks, even in oil-amended control treatments. When combined with its oligotrophic lifestyle (Suja et al., 2017), this suggests that *Colwellia* is well adapted to exploit transient, low-nutrient, early-stage oil inputs but is less competitive during later stages of oil weathering and nutrient-enriched conditions. Further, these findings suggest that *Colwellia* dominance is not universal following oil exposure, but instead depends on environmental context, spill response strategies, and microbial niche availability.

The dominance of *Pseudomonas* and *Pseudoalteromonas* was observed exclusively in biostimulated treatments, in which, after approximately 2.5 weeks, *Pseudomonas* populations increased markedly, coinciding with a decline in relative abundance of *Pseudoalteromonas* and other community members. Nutrient enrichment likely favored fast-growing, copiotrophic taxa with high metabolic flexibility, enabling *Pseudomonas* to outcompete other community members, as previously observed (Suja et al., 2017; Murphy et al., 2021). Although *Pseudoalteromonas* persisted, its relative decline suggests competitive exclusion as *Pseudomonas* populations expanded under nutrient-replete conditions. In contrast, control treatments lacking nutrient and dispersant amendments maintained a more diverse community structure and did not converge toward dominance by a limited number of taxa. *Pseudomonas* is a paradigmatic bacterial genus associated with both alkane and aromatic degradation (Mason et al., 2012; Pasumarthi et al., 2013; Nogales et al., 2017), whereas *Pseudoalteromonas* has been reported to persist in environments enriched in aromatic hydrocarbons (Dubinsky et al., 2013). Although direct interactions between these taxa were not examined in this study, the observed shift toward *Pseudomonas* dominance under nutrient-rich, biostimulated conditions is consistent with competitive dynamics favoring metabolically versatile alkane-degrading taxa. This pattern occurred despite the continued presence of both n-alkanes and PAHs in the system, suggesting that nutrient availability and substrate accessibility, rather than hydrocarbon presence alone, played a key role in structuring the microbial community.

*Sulfitobacter* is a common and often dominant member of surface-water microbial communities in Hudson Bay and other polar marine systems, where it participates in the transformation of inorganic sulfur compounds (Buchan et al., 2005). Consistent with this, *Sulfitobacter* was the most abundant taxon in the initial seawater before oil addition. In oil-only control treatments, *Sulfitobacter* relative abundance gradually declined over the six-week incubation period following diesel addition, while its relative abundance rapidly declined in biostimulated treatments, with almost no detection after 1 week. This rapid decline reflects strong environmental selectivity under nutrient- and dispersant-replenished conditions, in which *Sulfitobacter* may be at a competitive disadvantage.

Our results show that microbial community composition shifted rapidly within 1 to 2.5 weeks in response to nutrient and dispersant amendments, indicating that biostimulated conditions strongly and selectively favored hydrocarbon-degrading taxa. This rapid restructuring suggests that nutrient and dispersant additions accelerate microbial succession by enhancing hydrocarbon bioavailability and promoting the growth of metabolically versatile putative oil degraders. The Hudson Bay surface-water microbial community appears to include populations capable of responding to hydrocarbon inputs, with *Pseudomonas* and *Pseudoalteromonas* emerging as dominant taxa under both ambient and biostimulated conditions. Together, these findings indicate that while a diverse background community is present under natural conditions, biostimulation can rapidly select for a narrow subset of highly competitive hydrocarbon-degrading bacteria.

### 4.2 UV exposure causes greater loss of PAHs than of n-alkanes

UV-weathered Churchill diesel was used to simulate conditions conducive to MOS formation (Passow, 2016). Photochemically aged oil has been shown to promote marine snow formation, though the resulting aggregates are typically smaller and more compact than those formed with fresh oil (Passow, 2016). In our roller incubations, the MOS observed was relatively compact, which is consistent with these previous observations. This may be related to the higher viscosity and altered interfacial properties of irradiated diesel, which has lost volatile compounds and contains more surface-active photooxidation products (Janus et al., 2019). PAHs are generally more susceptible to photooxidation than n-alkanes (Bacosa et al., 2015), which can alter their environmental fate following an oil spill. Consistent with this, our UV-weathering experiments showed a greater loss of PAHs (22.2%) compared to n-alkanes (3.3%). Among the PAHs, pyrene, a high molecular weight compound, experienced the highest loss (68.2%) under UV exposure, reflecting its susceptibility to photooxidation. While photooxidation reduced PAH abundance, UV irradiation can also generate more reactive or toxic intermediates through photo-activation and photo-modification (Ekunwe et al., 2005), which may influence microbial degradation pathways and the potential toxicity of oil-contaminated waters. These changes in hydrocarbon composition could affect MOS formation and microbial community dynamics, as altered PAH and alkane profiles modify the chemical environment available for microbial colonization and hydrocarbon degradation.

### 4.3 MOS influences elevated partitioning of high molecular weight hydrocarbons

The observation of higher oil-to-aqueous ratios for n-alkanes in the control treatments suggests that, in the absence of dispersants or MOS formation, these compounds largely remain in the bulk oil phase. This is consistent with the hydrophobic nature of n-alkanes (Turner et al., 2014; Desmond et al., 2021), which limits their spontaneous partitioning into the aqueous phase. In biostimulated treatments, overall oil-to-water ratios were markedly lower, consistent with dispersant-mediated solubilization or incorporation into marine oil snow, followed by microbial processing of hydrocarbons. Therefore, the resulting higher oil:water ratios from lower aqueous concentrations in control treatments indicate limited natural dispersion and a reduced immediate substrate availability for hydrocarbonoclastic microbes, which may slow the early-stage biodegradation of these compounds compared to treatments with enhanced partitioning mechanisms.

LMW n-alkanes exhibited higher oil-to-water ratios compared to HMW n-alkanes in both treatments, which may reflect the relatively higher bioavailability and greater susceptibility of LMW compounds to microbial uptake and biodegradation. These observations highlight how both molecular weight and treatment regime influence hydrocarbon fate, with LMW compounds more readily mobilized and degraded in the aqueous phase than HMW homologues, as described by previous research (Singh et al., 2012).

Overall, lower oil-to-water ratios observed for PAHs compared to n-alkanes likely reflect their relatively higher aqueous solubility (Desmond et al., 2022) and structural characteristics (Zhang et al., 2015) that promote partial partitioning into the water phase. In contrast, the more hydrophobic n-alkanes remain largely in the bulk oil phase, leading to higher oil-to-water ratios in the absence of dispersants or MOS formation.

Biostimulated treatments exhibited markedly lower oil-to-water ratios for PAHs compared to controls, reflecting enhanced mobilization and biodegradation facilitated by dispersant action and incorporation into marine oil snow. LMW PAHs were more readily partitioned and degraded, whereas HMW PAHs were retained more in the water column, likely via MOS sorption, rather than being biodegraded, a pattern attributed to their lower aqueous solubility and greater incorporation into the organic fraction, which limits microbial access (Achten et al., 2011; Frapiccini and Marini, 2015). This interpretation is consistent with previous observations showing that dispersant addition promotes the incorporation and enrichment of HMW oil compounds within MOS (Wirth et al., 2018).

Phenanthrene and its alkylated homologues (1-methylphenanthrene and 2-methylphenanthrene) are frequently among the most abundant PAH compounds detected in petroleum-contaminated marine environments (Wang et al., 2011), making their transport behavior particularly relevant for assessing oil spill impacts. Moreover, surveys of monomethylated phenanthrene isomers have shown that these compounds can be two- to five-fold more toxic than parent phenanthrene (Lee and McCain, 2015), highlighting their ecological significance. The enhanced partitioning of these compounds under biostimulated conditions, therefore, suggests that MOS-mediated transport may increase the mobility and potential bioavailability of toxic PAHs to pelagic organisms.

Overall, n-alkanes exhibited a higher percentage of degradation than PAHs, consistent with their susceptibility to biodegradation (Prince et al., 2016; Stout and Wang, 2016). Nonetheless, alkane and PAH losses did not differ substantially between control and biostimulated treatments, suggesting that alkane and PAH biodegradation were not strongly limited by nutrient or dispersant availability under the conditions tested. This indicates that the microbial community sampled from Hudson Bay surface waters possessed an inherent capacity to efficiently degrade alkanes and PAHs, even in the absence of external stimulation. While biostimulation did not significantly increase total alkane and PAH degradation, it altered hydrocarbon partitioning dynamics, with increased transfer of HMW compounds into the aqueous phase through sorption via MOS formation. The absence of independent nutrient-only and dispersant-only treatments prevents differentiation between physicochemical dispersion effects and biologically mediated aggregation processes such as MOS formation. It is therefore unclear whether enhanced partitioning resulted primarily from dispersant-induced emulsification or microbially driven EPS production and aggregate formation. Future studies incorporating factorial treatment designs and direct quantification of MOS would help disentangle these mechanisms.

## 5 Conclusion

Findings of this study indicate that biostimulation through nutrient and dispersant addition substantially alters the fate and transport of diesel-derived hydrocarbons in Arctic marine surface waters by promoting marine oil snow formation and restructuring microbial communities with hydrocarbon-degrading taxa such as *Pseudomonas* and *Pseudoalteromonas* becoming dominant within weeks. Enhanced transfer of high-molecular-weight alkanes and selected PAHs into the aqueous phase under stimulated conditions suggests that MOS potentially acts as an effective vector for redistributing hydrophobic hydrocarbons within the water column. Although biostimulation can accelerate the microbial responses to oil contamination, it may modify the hydrocarbon transport pathways in ways that influence ecological risk. Understanding these trade-offs is critical for informing oil spill response strategies in cold and sensitive marine environments such as the Arctic.

## Data availability

All raw 16S rRNA gene sequence reads generated from this experiment have been deposited at NCBI’s Sequence Read Archive (SRA) under the BioProject PRJNA1176269 (accession numbers SAMN54986455 to SAMN54986446).

## Author contributions

**Thamali Vidanage** – Conceptualization, Data curation, Formal analysis, Methodology, Investigation, Visualization, Writing – original draft, Writing – review & editing, Project administration

**Durell Desmond** – Conceptualization, Methodology, Investigation, Data curation, Formal analysis, Validation, Writing – original draft, Writing – review & editing

**Dustin Isleifson** – Writing – review & editing, Funding acquisition

**Gary Stern** – Resources, Funding acquisition

**R. Eric Collins** – Conceptualization, Methodology, Validation, Resources, Writing – review & editing, Visualization, Supervision, Funding acquisition, Project administration

## Acknowledgements

We would like to express our sincere gratitude to David Binne and Debbie Armstrong for their support in acquiring water samples from the CMO-OSIM research facility. We also thank the technical staff at CEOS, including Jocelyn Plouffe, for sample preparation, and Lisa Oswald and Shiva Lashkari for their assistance with GC-MS analysis. Additionally, we thank Cyril Fredlund for his support during the fieldwork at the CMO.

## Supplementary material

### Additional figures

**Figure S1.**
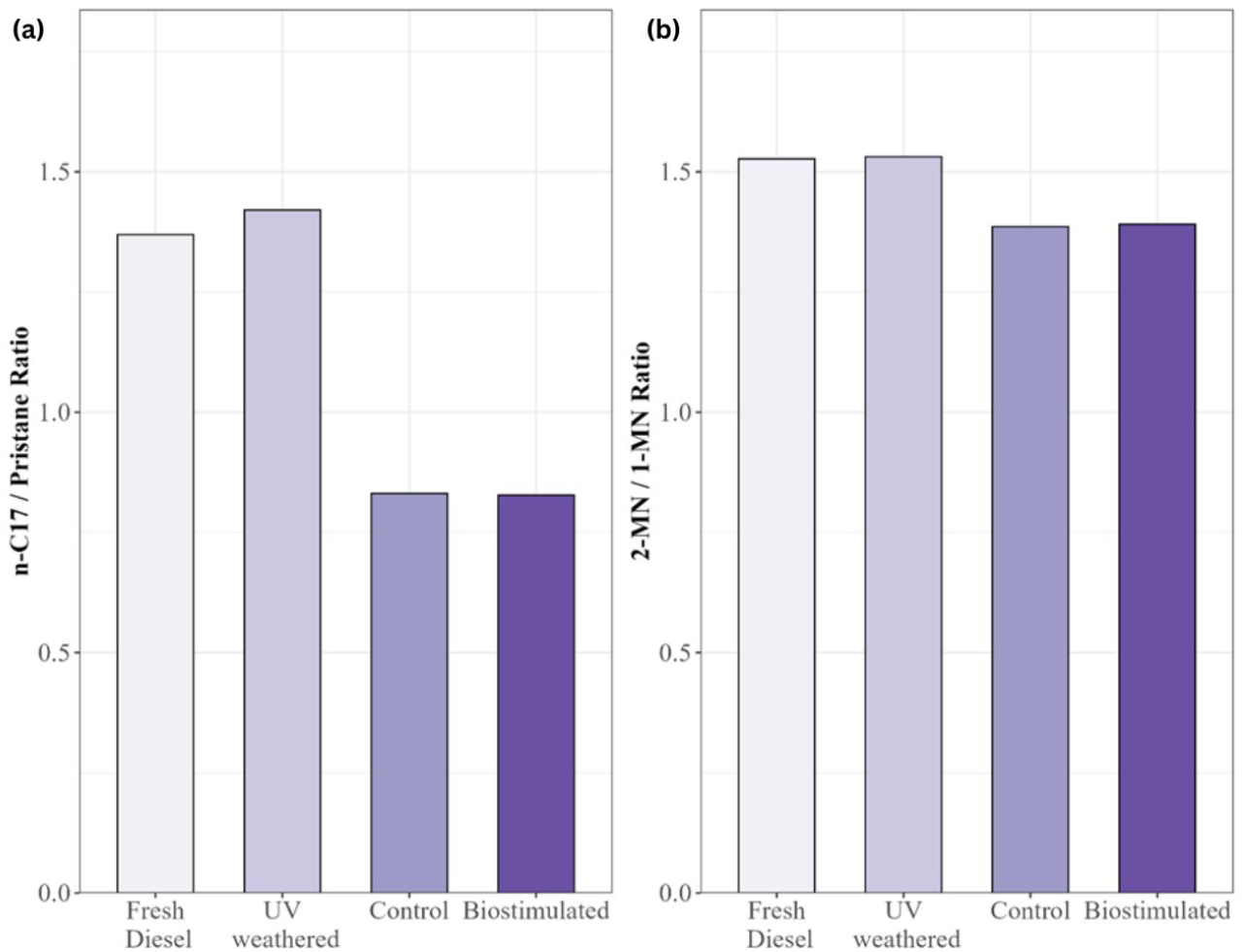
Diagnostic ratios for the biodegradation of n-alkanes (a) and PAHs (b) after 6 weeks of incubation.

**Figure S2.**
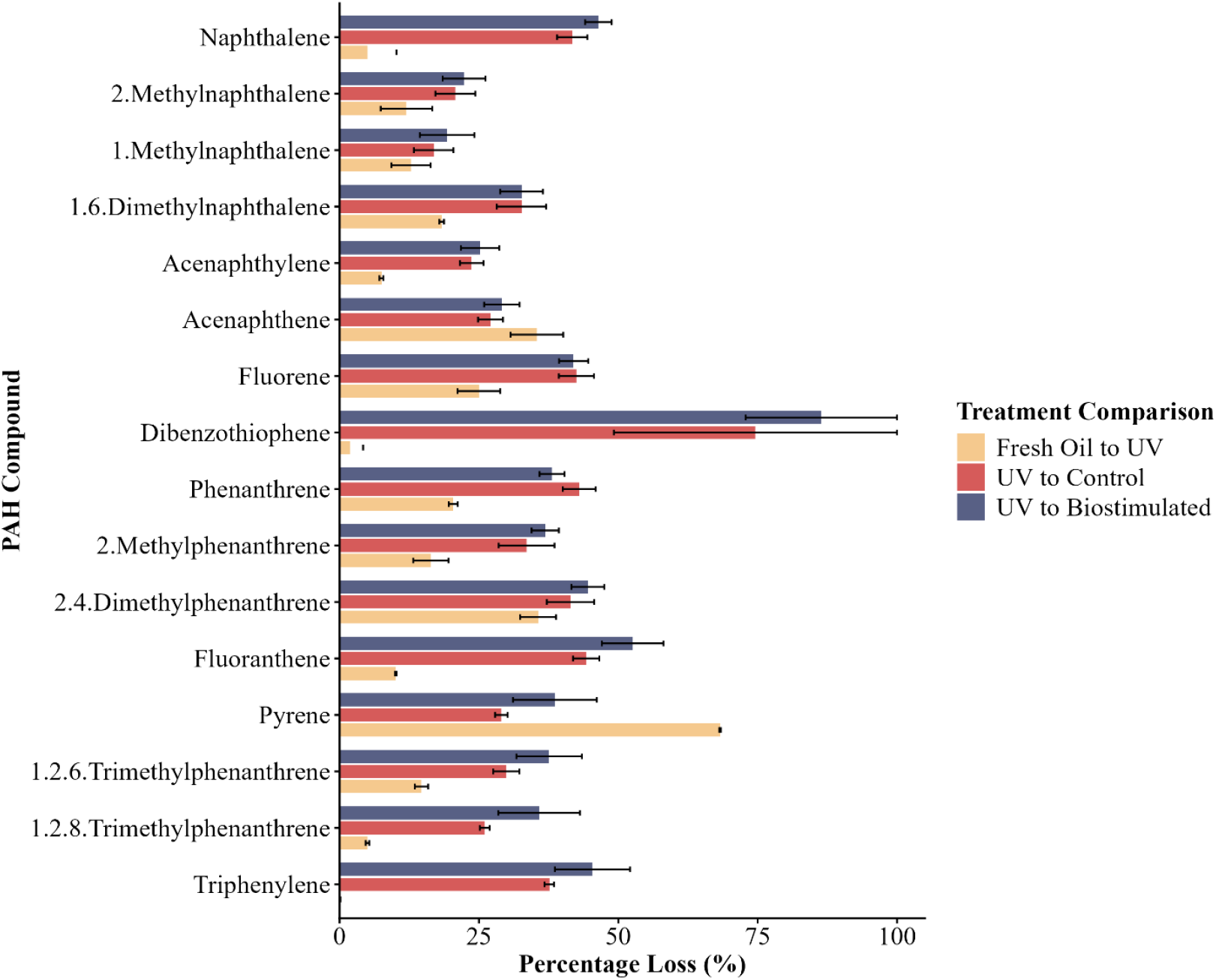
Percentage loss of individual PAH compounds during the 6-week incubations. Bars represent the mean percentage loss (%) for each compound. Error bars indicate the standard error of the mean (SE). Compounds are arranged in order of increasing molecular weight. The data are based on the bulk oil phase.

**Figure S3.**
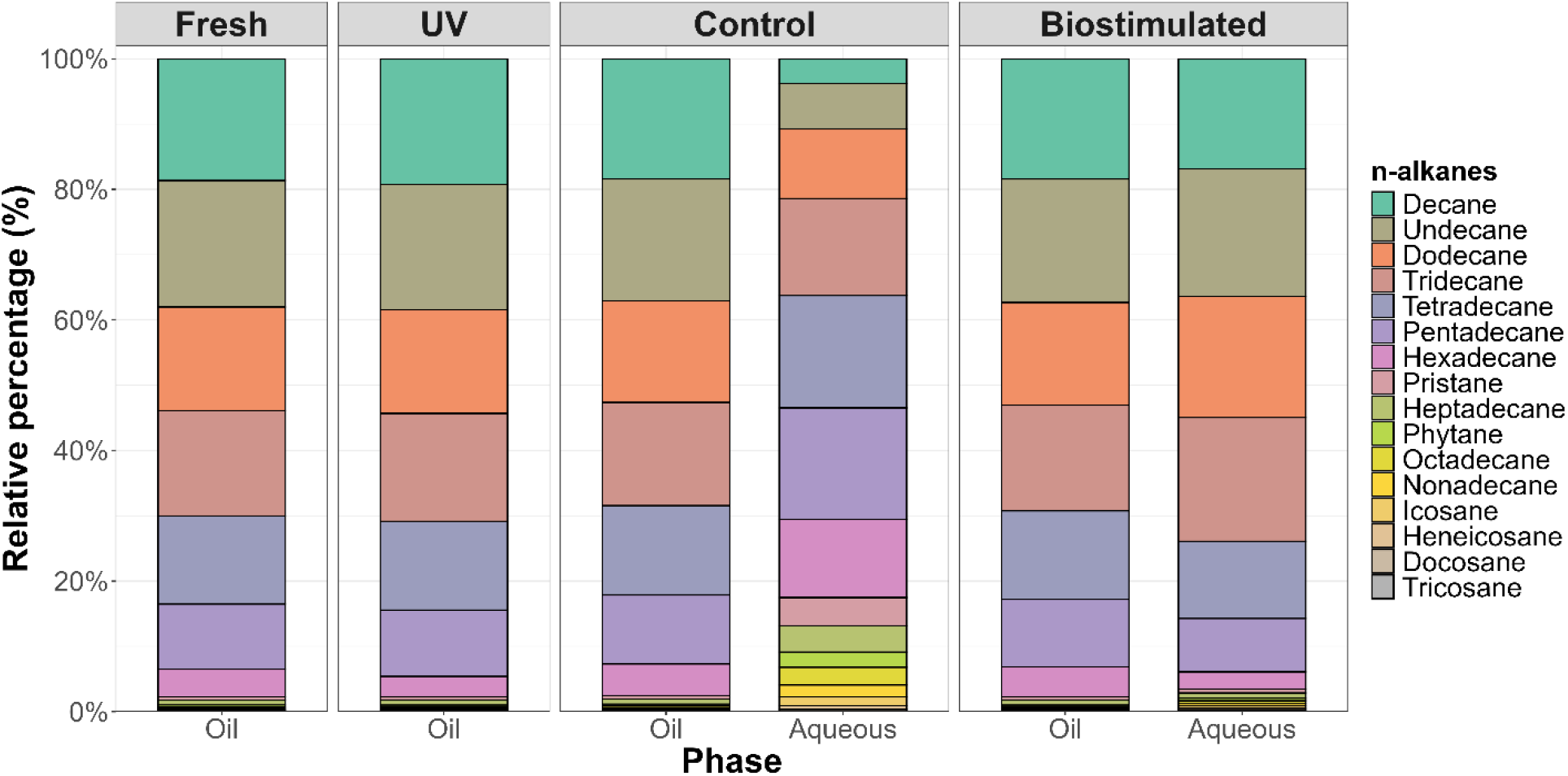
Relative concentration (%) of individual n-alkanes in fresh and UV-weathered diesel, and in the control and biostimulated treatments after incubation. For control and biostimulated treatments, n-alkane distributions are shown separately for the oil and aqueous phases. Values represent the percentage contribution of each compound to the total measured n-alkane concentration of each phase within each sample.

**Figure S4.**
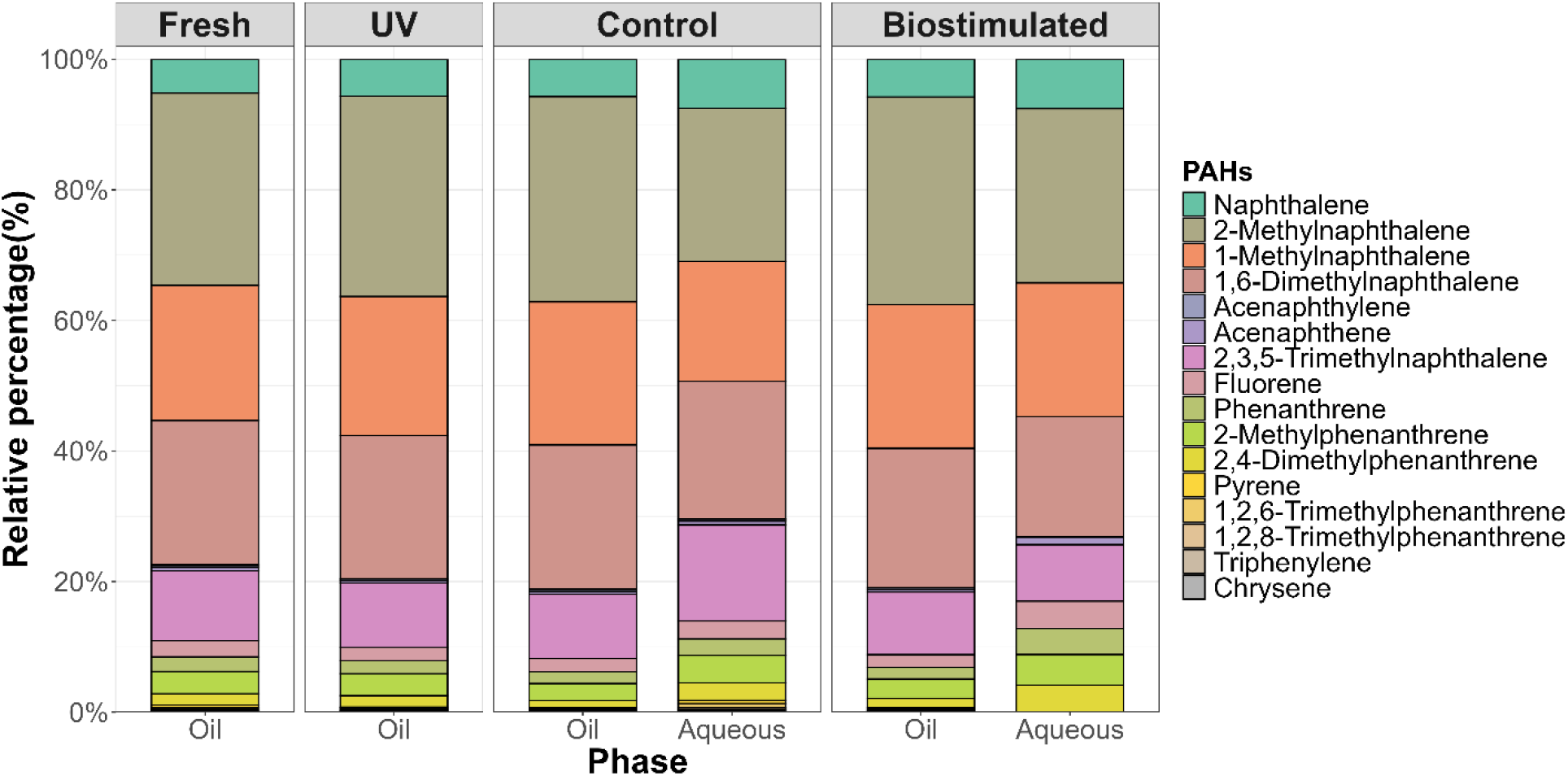
Relative concentration (%) of individual PAHs in fresh and UV-weathered diesel, and in the control and biostimulated treatments after incubation. For control and biostimulated treatments, PAH distributions are shown separately for the oil and aqueous phases. Values represent the percentage contribution of each compound to the total measured PAH concentration of each phase within each sample.

### GC-MS instrumental parameters

**Table S1.** Table of identified compounds and their multiple reaction monitoring mass spectrometry method quantifier settings.

| Investigated Compounds | Standard Quantifiers | Precursor Ion | Product Ion | Dwell | CE (eV) |
|---|---|---|---|---|---|
| Naphthalene | Naphthalene | 128 | 128 | 54.7 | 5 |
| 2-Methylnaphthalene | 2-Methylnaphthalene | 142 | 142 | 54.8 | 5 |
| 1-Methylnaphthalene | 1-Methylnaphthalene | 142 | 142 | 54.8 | 5 |
| 1,6-Dimethylnaphthalene | 1,6-Dimethylnaphthalene | 156 | 156 | 110.3 | 5 |
| Acenaphthylene | Acenaphthylene | 152 | 152 | 32.5 | 5 |
| Acenaphthene | Acenaphthene | 154 | 154 | 32.5 | 5 |
| 2,3,5-Trimethylnaphthalene | 2,3,5-Trimethylnaphthalene | 170 | 170 | 110.4 | 5 |
| Fluorene | Fluorene | 166 | 165 | 40.9 | 26 |
| Dibenzothiophene | Dibenzothiophene | 184 | 184 | 110.3 | 5 |
| Phenanthrene | Phenanthrene | 178 | 178 | 40.9 | 5 |
| Anthracene | Anthracene | 178 | 178 | 40.9 | 5 |
| 2-Methyldibenzothiophene | 2-Methyldibenzothiophene | 198 | 198 | 54.8 | 5 |
| 2-Methylphenanthrene | 2-Methylphenanthrene | 192 | 192 | 54.8 | 5 |
| 2,8-Dimethyldibenzothiophene | 2,8-Dimethyldibenzothiophene | 212 | 212 | 110.4 | 5 |
| 2,4-Dimethylphenanthrene | 2,4-Dimethylphenanthrene | 206 | 206 | 54.8 | 5 |
| 2,4,7-Trimethyldibenzothiophene | 2,4,7-Trimethyldibenzothiophene | 226 | 226 | 54.8 | 5 |
| Fluoranthene | Fluoranthene | 202 | 202 | 54.8 | 5 |
| Pyrene | Pyrene | 202 | 202 | 41 | 5 |
| 1,2,6-Trimethylphenanthrene | 1,2,6-Trimethylphenanthrene | 220 | 220 | 36.3 | 5 |
| 1,2,8-Trimethylphenanthrene | 1,2,8-Trimethylphenanthrene | 220 | 220 | 36.3 | 5 |
| 1-Methylfluoranthene | 1-Methylfluoranthene | 216 | 216 | 36.3 | 10 |
| Benzo[e]phenanthrene | Benzo[e]phenanthrene | 228 | 228 | 110.4 | 5 |
| Benz[a]anthracene | Benz[a]anthracene | 228 | 228 | 36.2 | 14 |
| Triphenylene | Triphenylene | 228 | 228 | 46.8 | 5 |
| Chrysene | Chrysene | 228 | 228 | 46.8 | 5 |
| 1-Methylchrysene | 1-Methylchrysene | 242 | 242 | 54.8 | 5 |
| 6-Ethylchrysene | 6-Ethylchrysene | 256 | 241 | 54.8 | 18 |
| Benzo[b]fluoranthene | Benzo[b]fluoranthene | 252 | 250 | 21.4 | 50 |
| Benzo[k]fluoranthene | Benzo[k]fluoranthene | 252 | 252 | 2.14 | 14 |
| Benzo[j]fluoranthene | Benzo[j]fluoranthene | 252 | 252 | 21.4 | 14 |
| Benzo[e]pyrene | Benzo[e]pyrene | 252 | 252 | 36.2 | 14 |
| Benzo[a]pyrene | Benzo[a]pyrene | 252 | 252 | 36.2 | 14 |
| Perylene | Perylene | 252 | 252 | 54.7 | 14 |
| Indeno[1,2,3-c,d]pyrene | Indeno[1,2,3-c,d]pyrene | 276 | 276 | 32.5 | 14 |
| Dibenz[a,h]anthracene | Dibenz[a,h]anthracene | 278 | 278 | 32.5 | 14 |
| Benzo[g,h,i]perylene | Benzo[g,h,i]perylene | 276 | 276 | 54.8 | 14 |
| Decane | Decane | 57 | 57 | 110.2 | 0 |
| Undecane | Undecane | 57 | 57 | 110.2 | 0 |
| Dodecane | Dodecane | 57 | 57 | 54.7 | 0 |
| Tridecane | Tridecane | 57 | 57 | 110.2 | 0 |
| Tetradecane | Tetradecane | 57 | 57 | 100.2 | 0 |
| Pentadecane | Pentadecane | 57 | 57 | 110.2 | 0 |
| Hexadecane | Hexadecane | 57 | 57 | 36.2 | 0 |
| Pristane | Pristane | 57 | 57 | 54.8 | 0 |
| Heptadecane | Heptadecane | 57 | 57 | 54.8 | 0 |
| Phytane | Phytane | 57 | 57 | 54.8 | 0 |
| Octadecane | Octadecane | 57 | 57 | 54.8 | 0 |
| Nonadecane | Nonadecane | 57 | 57 | 110.2 | 0 |
| Eicosane | Eicosane | 57 | 57 | 54.7 | 0 |
| Heneicosane | Heneicosane | 57 | 57 | 110.2 | 0 |
| Docosane | Docosane | 57 | 57 | 110.2 | 0 |
| Tricosane | Tricosane | 57 | 57 | 110.2 | 0 |
| Tetracosane | Tetracosane | 71 | 71 | 40.8 | 0 |
| Pentacosane | Pentacosane | 57 | 57 | 110.2 | 0 |
| Hexacosane | Hexacosane | 57 | 57 | 110.2 | 0 |
| Heptacosane | Heptacosane | 57 | 57 | 110.2 | 0 |
| Octacosane | Octacosane | 57 | 57 | 54.7 | 0 |
| Nonacosane | Nonacosane | 57 | 57 | 110.2 | 0 |
| Triacontane | Triacontane | 57 | 57 | 110.2 | 0 |
| Hentriacontane | Hentriacontane | 57 | 57 | 54.7 | 0 |
| Dotriacontane | Dotriacontane | 57 | 57 | 110.2 | 0 |
| Tritriacontane | Tritriacontane | 57 | 57 | 110.2 | 0 |
| Tetratriacontane | Tetratriacontane | 57 | 57 | 110.2 | 0 |
| Pentatriacontane | Pentatriacontane | 57 | 57 | 54.7 | 0 |
| Hexatriacontane | Hexatriacontane | 57 | 57 | 110.2 | 0 |
| Heptatriacontane | Heptatriacontane | 57 | 57 | 110.2 | 0 |
| Octatriacontane | Octatriacontane | 57 | 57 | 110.2 | 0 |
| Nonatriacontane | Nonatriacontane | 57 | 57 | 110.2 | 0 |
| Tetracontane | Tetracontane | 57 | 57 | 110.2 | 0 |

## References

Christine Achten, Shubo Cheng, Kristina L. Straub, and Thilo Hofmann. The lack of microbial degradation of polycyclic aromatic hydrocarbons from coal-rich soils. Environmental Pollution, 159(2):623–629, 2011. doi: 10.1016/j.envpol.2010.09.035.

Saidat Omowunmi Adeniran-Obey and David Agbontaen Osagie. Impacts of Invasive Species on the Arctic Environment. In P. O. Isibor, editor, Arctic Marine Ecotoxicology: Climate Change, Pollutants, and Their Far-Reaching Effects, pages 381–402. Springer Nature Switzerland, Cham, 2024. doi: 10.1007/978-3-031-73584-4_18.

Jackie Aislabie, David J. Saul, and Julia M. Foght. Bioremediation of hydrocarbon-contaminated polar soils. Extremophiles, 10(3):171–179, 2006. doi: 10.1007/s00792-005-0498-4.

Jonathan Andrews, David Babb, and David G. Barber. Climate change and sea ice: Shipping accessibility on the marine transportation corridor through Hudson Bay and Hudson Strait (1980–2014). Elementa: Science of the Anthropocene, 5:15, 2017. doi: 10.1525/elementa.130.

Jonathan Andrews, David Babb, and David G. Barber. Climate change and sea ice: Shipping in Hudson Bay, Hudson Strait, and Foxe Basin (1980–2016). Elementa: Science of the Anthropocene, 6:19, 2018. doi: 10.1525/elementa.281.

Carol Arnosti, Kai Ziervogel, Tingting Yang, and Andreas Teske. Oil-derived marine aggregates – hot spots of polysaccharide degradation by specialized bacterial communities. Deep Sea Research Part II: Topical Studies in Oceanography, 129:179–186, 2016. doi: 10.1016/j.dsr2.2014.12.008.

Jasmin C. Athas, Kelly Jun, Caitlyn McCafferty, Olasehinde Owoseni, Vijay T. John, and Srinivasa R. Raghavan. An Effective Dispersant for Oil Spills Based on Food-Grade Amphiphiles. Langmuir, 30(31): 9285–9294, 2014. doi: 10.1021/la502312n.

Ronald M. Atlas and Terry C. Hazen. Oil Biodegradation and Bioremediation: A Tale of the Two Worst Spills in U.S. History. Environmental Science & Technology, 45(16):6709–6715, 2011. doi: 10.1021/es2013227.

Hernando P. Bacosa, Deana L. Erdner, and Zhanfei Liu. Differentiating the roles of photooxidation and biodegradation in the weathering of Light Louisiana Sweet crude oil in surface water from the Deepwater Horizon site. Marine Pollution Bulletin, 95(1):265–272, 2015. doi: 10.1016/j.marpolbul.2015.04.005.

Huifang Bi, Zheng Wang, Rengyu Yue, Jiyao Sui, Catherine N. Mulligan, Kenneth Lee, Scott Pegau, Zhi Chen, and Chunjiang An. Oil spills in coastal regions of the Arctic and Subarctic: Environmental impacts, response tactics, and preparedness. Science of The Total Environment, 958:178025, 2025. doi: 10.1016/j.scitotenv.2024.178025.

Alison Buchan, José M. González, and Mary Ann Moran. Overview of the Marine *Roseobacter* Lineage. Applied and Environmental Microbiology, 71(10):5665–5677, 2005. doi: 10.1128/aem.71.10.5665-5677. 2005.

Benjamin J Callahan, Paul J McMurdie, Michael J Rosen, Andrew W Han, Amy Jo A Johnson, and Susan P Holmes. DADA2: High-resolution sample inference from Illumina amplicon data. Nature Methods, 13(7): 581–583, 2016. doi: 10.1038/nmeth.3869.

Jackie Dawson, Natalie Carter, Nicolien van Luijk, Colleen Parker, Melissa Weber, Alison Cook, Kayla Grey, and Jennifer Provencher. Infusing Inuit and local knowledge into the Low Impact Shipping Corridors: An adaptation to increased shipping activity and climate change in Arctic Canada. Environmental Science & Policy, 105:19–36, 2020. doi: 10.1016/j.envsci.2019.11.013.

Department of Finance Canada. Budget. https://budget.canada.ca/2025/report-rapport/overview-apercu-en.html#a5, 2025.

Durell S. Desmond, Diana Saltymakova, Alastair Smith, Teresinha Wolfe, Nolan Snyder, Katarzyna Polcwiartek, María Bautista, Marcos Lemes, Casey R. J. Hubert, David G. Barber, Dustin Isleifson, and Gary A. Stern. Photooxidation and biodegradation potential of a light crude oil in first-year sea ice. Marine Pollution Bulletin, 165:112154, 2021. doi: 10.1016/j.marpolbul.2021.112154.

Durell S. Desmond, Diana Saltymakova, Odile Crabeck, Georg Schreckenbach, James D. Xidos, David G. Barber, Dustin Isleifson, and Gary A. Stern. Methods for Interpreting the Partitioning and Fate of Petroleum Hydrocarbons in a Sea Ice Environment. The Journal of Physical Chemistry A, 126(5):772–786, 2022. doi: 10.1021/acs.jpca.1c08357.

Eric A. Dubinsky, Mark E. Conrad, Romy Chakraborty, Markus Bill, Sharon E. Borglin, James T. Hollibaugh, Olivia U. Mason, Yvette M. Piceno, Francine C. Reid, William T. Stringfellow, Lauren M. Tom, Terry C. Hazen, and Gary L. Andersen. Succession of Hydrocarbon-Degrading Bacteria in the Aftermath of the Deepwater Horizon Oil Spill in the Gulf of Mexico. Environmental Science & Technology, 47(19): 10860–10867, 2013. doi: 10.1021/es401676y.

Victor M. Eguíluz, Juan Fernández-Gracia, Xabier Irigoien, and Carlos M. Duarte. A quantitative assessment of Arctic shipping in 2010–2014. Scientific Reports, 6(1):30682, 2016. doi: 10.1038/srep30682.

Stephen I. N. Ekunwe, Rochelle D. Hunter, and Huey-Min Hwang. Ultraviolet Radiation Increases the Toxicity of Pyrene, 1-Aminopyrene and 1-Hydroxypyrene to Human Keratinocytes. International Journal of Environmental Research and Public Health, 2(1):58–62, 2005. doi: 10.3390/ijerph2005010058.

Emanuela Frapiccini and Mauro Marini. Polycyclic Aromatic Hydrocarbon Degradation and Sorption Parameters in Coastal and Open-Sea Sediment. Water, Air, & Soil Pollution, 226(8):246, 2015. doi: 10.1007/s11270-015-2510-7.

C. L. Friedman, J. R. Pierce, and N. E. Selin. Assessing the Influence of Secondary Organic versus Primary Carbonaceous Aerosols on Long-Range Atmospheric Polycyclic Aromatic Hydrocarbon Transport. Environmental Science & Technology, 48(6):3293–3302, 2014. doi: 10.1021/es405219r.

Jie Fu, Yanyan Gong, Xiao Zhao, S. E. O’Reilly, and Dongye Zhao. Effects of Oil and Dispersant on Formation of Marine Oil Snow and Transport of Oil Hydrocarbons. Environmental Science & Technology, 48(24):14392–14399, 2014. doi: 10.1021/es5042157.

S. C. Gad. Diesel Fuel. In Encyclopedia of Toxicology (Third Edition). Elsevier, 2014. doi: 10.1016/B978-0-12-386454-3.00837-X.

Marie-Ève Garneau, Christine Michel, Guillaume Meisterhans, Nathalie Fortin, Thomas L. King, Charles W. Greer, and Kenneth Lee. Hydrocarbon biodegradation by Arctic sea-ice and sub-ice microbial communities during microcosm experiments, Northwest Passage (Nunavut, Canada). FEMS Microbiology Ecology, 92 (10):fiw130, 2016. doi: 10.1093/femsec/fiw130.

Benjamin H. Gregson, Boyd A. McKew, Robert D. Holland, Timothy J. Nedwed, Roger C. Prince, and Terry J. McGenity. Marine Oil Snow, a Microbial Perspective. Frontiers in Marine Science, 8:619484, 2021. doi: 10.3389/fmars.2021.619484.

Tony Gutierrez, David Berry, Tingting Yang, Sara Mishamandani, Luke McKay, Andreas Teske, and Michael D. Aitken. Role of Bacterial Exopolysaccharides (EPS) in the Fate of the Oil Released during the Deepwater Horizon Oil Spill. PLoS ONE, 8(6):e67717, 2013. doi: 10.1371/journal.pone.0067717.

Esteban Góngora, Ya-Jou Chen, Madison Ellis, Mira Okshevsky, and Lyle Whyte. Hydrocarbon bioremediation on Arctic shorelines: Historic perspective and roadway to the future. Environmental Pollution, 305: 119247, 2022. doi: 10.1016/j.envpol.2022.119247.

Terry C. Hazen, Roger C. Prince, and Nagissa Mahmoudi. Marine Oil Biodegradation. Environmental Science & Technology, 50(5):2121–2129, 2016. doi: 10.1021/acs.est.5b03333.

Rafał Janus, Karol Kołomanski, Mariusz Wadrzyk, and Marek Lewandowski. Degradation of petroleum diesel fuel accelerated by UV irradiation: the impact of ageing on chemical composition and selected physicochemical properties. E3S Web of Conferences, 108:02003, 2019. doi: 10.1051/e3sconf/201910802003.

Colin D. Johnston, Trevor P. Bastow, and Nathan L. Innes. The use of biodegradation signatures and biomarkers to differentiate spills of petroleum hydrocarbon liquids in the subsurface and estimate natural mass loss. European Journal of Soil Biology, 43(5-6):328–334, 2007. doi: 10.1016/j.ejsobi.2007.03.008.

B. Judson. Trends In Canadian Arctic Shipping Traffic – Myths And Rumours. OnePetro, 2010. Accessed January 30, 2026.

F. Lasserre. Canadian arctic marine transportation issues, opportunities and challenges. doi: 10.11575/sppp.v15i1.72626, 2022.

James S. Latimer and Jinshu Zheng. The Sources, Transport, and Fate of PAHs in the Marine Environment. In P. E. T. Douben, editor, PAHs: An Ecotoxicological Perspective, pages 7–33. Wiley, 2003. doi: 10.1002/0470867132.ch2.

J G Leahy and R R Colwell. Microbial degradation of hydrocarbons in the environment. Microbiological Reviews, 54(3):305–315, 1990. doi: 10.1128/mr.54.3.305-315.1990.

Sang Ho Lee and Wilfred C. McCain. Wildlife Toxicity Assessment for Phenanthrene. In Wildlife Toxicity Assessments for Chemicals of Military Concern, pages 591–598. Elsevier, 2015. doi: 10.1016/b978-0-12-800020-5.00031-4.

Fuyan Liang, Mingming Lu, Tim C. Keener, Zifei Liu, and Soon-Jai Khang. The organic composition of diesel particulate matter, diesel fuel and engine oil of a non-road diesel generator. Journal of Environmental Monitoring, 7(10):983–988, 2005. doi: 10.1039/b504728e.

Marcel Martin. Cutadapt removes adapter sequences from high-throughput sequencing reads. EMBnet.journal, 17(1):10–12, 2011. doi: 10.14806/ej.17.1.200.

Olivia U Mason, Terry C Hazen, Sharon Borglin, Patrick S G Chain, Eric A Dubinsky, Julian L Fortney, James Han, Hoi-Ying N Holman, Jenni Hultman, Regina Lamendella, Rachel Mackelprang, Stephanie Malfatti, Lauren M Tom, Susannah G Tringe, Tanja Woyke, Jizhong Zhou, Edward M Rubin, and Janet K Jansson. Metagenome, metatranscriptome and single-cell sequencing reveal microbial response to Deepwater Horizon oil spill. The ISME Journal, 6(9):1715–1727, 2012. doi: 10.1038/ismej.2012.59.

Olivia U. Mason, James Han, Tanja Woyke, and Janet K. Jansson. Single-cell genomics reveals features of a *Colwellia* species that was dominant during the Deepwater Horizon oil spill. Frontiers in Microbiology, 5, 2014. doi: 10.3389/fmicb.2014.00332.

Lawrence R. Mudryk, Jackie Dawson, Stephen E. L. Howell, Chris Derksen, Thomas A. Zagon, and Mike Brady. Impact of 1, 2 and 4 °C of global warming on ship navigation in the Canadian Arctic. Nature Climate Change, 11(8):673–679, 2021. doi: 10.1038/s41558-021-01087-6.

Albertinka J. Murk, David J. Hollander, Shuangling Chen, Chuanmin Hu, Yongxue Liu, Sophie M. Vonk, Patrick T. Schwing, Sherryl Gilbert, and Edwin M. Foekema. A Predictive Strategy for Mapping Locations Where Future MOSSFA Events Are Expected. In S. A. Murawski, C. H. Ainsworth, S. Gilbert, D. J. Hollander, C. B. Paris, M. Schlüter, et al., editors, Scenarios and Responses to Future Deep Oil Spills: Fighting the Next War, pages 355–368. Springer International Publishing, Cham, 2020. doi: 10.1007/978-3-030-12963-7_21.

Sean M. C. Murphy, María A. Bautista, Margaret A. Cramm, and Casey R. J. Hubert. Diesel and Crude Oil Biodegradation by Cold-Adapted Microbial Communities in the Labrador Sea. Applied and Environmental Microbiology, 87(20):e00800–21, 2021. doi: 10.1128/aem.00800-21.

C. S. Neethu, C. Saravanakumar, R. Purvaja, R. S. Robin, and R. Ramesh. Oil-Spill Triggered Shift in Indigenous Microbial Structure and Functional Dynamics in Different Marine Environmental Matrices. Scientific Reports, 9(1):1354, 2019. doi: 10.1038/s41598-018-37903-x.

J. Nogales, J. L. García, and E. Díaz. Degradation of Aromatic Compounds in Pseudomonas: A Systems Biology View. In Aerobic Utilization of Hydrocarbons, Oils and Lipids, pages 1–49. Springer, Cham, 2017. doi: 10.1007/978-3-319-39782-5_32-1.

Julia Olsen, Natalie Ann Carter, and Jackie Dawson. Community perspectives on the environmental impacts of Arctic shipping: Case studies from Russia, Norway and Canada. Cogent Social Sciences, 5(1):1609189, 2019. doi: 10.1080/23311886.2019.1609189.

PAME. The increase in Arctic Shipping: 2013-2024. Protection of the Arctic Marine Environment (PAME), 2024. URL https://hdl.handle.net/11374/2733.3. Accessed January 29, 2026.

U Passow, K Ziervogel, V Asper, and A Diercks. Marine snow formation in the aftermath of the Deepwater Horizon oil spill in the Gulf of Mexico. Environmental Research Letters, 7(3):035301, 2012. doi: 10.1088/1748-9326/7/3/035301.

Uta Passow. Formation of rapidly-sinking, oil-associated marine snow. Deep Sea Research Part II: Topical Studies in Oceanography, 129:232–240, 2016. doi: 10.1016/j.dsr2.2014.10.001.

Rajesh Pasumarthi, Sivaraman Chandrasekaran, and Srikanth Mutnuri. Biodegradation of crude oil by *Pseudomonas aeruginosa* and *Escherichia fergusonii* isolated from the Goan coast. Marine Pollution Bulletin, 76(1-2):276–282, 2013. doi: 10.1016/j.marpolbul.2013.08.026.

Angela Peeb, Nga Phuong Dang, Marika Truu, Hiie Nõlvak, Chris Petrich, and Jaak Truu. Assessment of Hydrocarbon Degradation Potential in Microbial Communities in Arctic Sea Ice. Microorganisms, 10(2): 328, 2022. doi: 10.3390/microorganisms10020328.

Larissa Pizzolato, Stephen E. L. Howell, Jackie Dawson, Frédéric Laliberté, and Luke Copland. The influence of declining sea ice on shipping activity in the Canadian Arctic. Geophysical Research Letters, 43(23): 12,146–12,154, 2016. doi: 10.1002/2016gl071489.

Roger C. Prince, Thomas S. Coolbaugh, and Thomas F. Parkerton. Oil dispersants do facilitate biodegradation of spilled oil. Proceedings of the National Academy of Sciences, 113(11):E1421–E1421, 2016. doi: 10.1073/pnas.1525333113.

Christian Quast, Elmar Pruesse, Pelin Yilmaz, Jan Gerken, Timmy Schweer, Pablo Yarza, Jörg Peplies, and Frank Oliver Glöckner. The SILVA ribosomal RNA gene database project: improved data processing and web-based tools. Nucleic Acids Research, 41(D1):D590–D596, 2013. doi: 10.1093/nar/gks1219.

Antonietta Quigg, Uta Passow, Kendra L. Daly, Adrian Burd, David J. Hollander, Patrick T. Schwing, and Kenneth Lee. Marine Oil Snow Sedimentation and Flocculent Accumulation (MOSSFA) Events: Learning from the Past to Predict the Future. In S. A. Murawski, C. H. Ainsworth, S. Gilbert, D. J. Hollander, C. B. Paris, M. Schlüter, et al., editors, Deep Oil Spills: Facts, Fate, and Effects, pages 196–220. Springer International Publishing, Cham, 2020. doi: 10.1007/978-3-030-11605-7_12.

Christopher M. Reddy, J. Samuel Arey, Jeffrey S. Seewald, Sean P. Sylva, Karin L. Lemkau, Robert K. Nelson, Catherine A. Carmichael, Cameron P. McIntyre, Judith Fenwick, G. Todd Ventura, Benjamin A. S. Van Mooy, and Richard Camilli. Composition and fate of gas and oil released to the water column during the Deepwater Horizon oil spill. Proceedings of the National Academy of Sciences, 109(50):20229–20234, 2012. doi: 10.1073/pnas.1101242108.

Deni Ribicic, Kelly Marie McFarlin, Roman Netzer, Odd Gunnar Brakstad, Anika Winkler, Mimmi Throne-Holst, and Trond Røvik Størseth. Oil type and temperature dependent biodegradation dynamics - Combining chemical and microbial community data through multivariate analysis. BMC Microbiology, 18(1):83, 2018. doi: 10.1186/s12866-018-1221-9.

Jesse Ross, David Hollander, Susan Saupe, Adrian B. Burd, Sherryl Gilbert, and Antonietta Quigg. Integrating marine oil snow and MOSSFA into oil spill response and damage assessment. Marine Pollution Bulletin, 165:112025, 2021. doi: 10.1016/j.marpolbul.2021.112025.

Aikaterini Sakellari, Sotirios Karavoltsos, Ipek Moutafis, Konstantinos Koukoulakis, Manos Dassenakis, and Evangelos Bakeas. Occurrence and distribution of polycyclic aromatic hydrocarbons in the marine surface microlayer of an industrialized coastal area in the eastern mediterranean. Water, 13(22):3174, 2021. doi: 10.3390/w13223174.

Katrine Scheibye, Jan H. Christensen, and Anders R. Johnsen. Biodegradation of crude oil in Arctic subsurface water from the Disko Bay (Greenland) is limited. Environmental Pollution, 223:73–80, 2017. doi: 10.1016/j.envpol.2016.12.032.

S. N. Singh, B. Kumari, and Shweta Mishra. Microbial Degradation of Alkanes. In S. N. Singh, editor, Microbial Degradation of Xenobiotics, pages 439–469. Springer, Berlin, Heidelberg, 2012. doi: 10.1007/978-3-642-23789-8_17.

C. Soriot, J. Stroeve, and A. Crawford. Record Early Sea Ice Loss in Southeastern Hudson Bay in Spring 2024. Geophysical Research Letters, 52(4):e2024GL112584, 2025. doi: 10.1029/2024gl112584.

Gary A. Stern, Colin R. Macdonald, Paloma Calabria Carvalho, Teresinha Wolfe, and Fernanda Ferraz. Baseline levels and characterization of hydrocarbons in surface marine sediments along the transportation corridor in Hudson Bay: A multivariate analysis of *n*-alkanes, PAHs and biomarkers. Science of The Total Environment, 855:158718, 2023. doi: 10.1016/j.scitotenv.2022.158718.

S. Stout and Z. Wang, editors. Standard Handbook Oil Spill Environmental Forensics: Fingerprinting and Source Identification. Academic Press, 2016.

Julienne Stroeve, Alex Crawford, Steve Ferguson, Ian Stirling, Louise Archer, Geoffrey York, David Babb, and Robbie Mallett. Ice-free period too long for Southern and Western Hudson Bay polar bear populations if global warming exceeds 1.6 to 2.6 °C. Communications Earth & Environment, 5(1):296, 2024. doi: 10.1038/s43247-024-01430-7.

Laura Duran Suja, Stephen Summers, and Tony Gutierrez. Role of EPS, Dispersant and Nutrients on the Microbial Response and MOS Formation in the Subarctic Northeast Atlantic. Frontiers in Microbiology, 8: 676, 2017. doi: 10.3389/fmicb.2017.00676.

Matthew Tarr, Pheobe Zito, Edward Overton, Gregory Olson, Puspa Adkikari, and Christopher Reddy. Weathering of Oil Spilled in the Marine Environment. Oceanography, 29(3):126–135, 2016. doi: 10.5670/oceanog.2016.77.

R. E. Turner, E. B. Overton, B. M. Meyer, M. S. Miles, and L. Hooper-Bui. Changes in the concentration and relative abundance of alkanes and PAHs from the Deepwater Horizon oiling of coastal marshes. Marine Pollution Bulletin, 86(1-2):291–297, 2014. doi: 10.1016/j.marpolbul.2014.07.003.

USEPA. Priority Pollutant List, 2014. URL https://www.epa.gov/sites/default/files/2015-09/documents/priority-pollutant-list-epa.pdf.

Justine S. van Eenennaam, Shokouh Rahsepar, Jagoš R. Radovic, Thomas B. P. Oldenburg, Jessica Wonink, Alette A. M. Langenhoff, Albertinka J. Murk, and Edwin M. Foekema. Marine snow increases the adverse effects of oil on benthic invertebrates. Marine Pollution Bulletin, 126:339–348, 2018. doi: 10.1016/j.marpolbul.2017.11.028.

Cuiping Wang, Hongwen Sun, Ying Chang, Zhiguang Song, and Xuebo Qin. PAHs distribution in sediments associated with gas hydrate and oil seepage from the Gulf of Mexico. Marine Pollution Bulletin, 62(12): 2714–2723, 2011. doi: 10.1016/j.marpolbul.2011.09.016.

H. Wickham. ggplot2: Elegant Graphics for Data Analysis. Springer-Verlag New York, 2016. URL https://ggplot2.tidyverse.org.

Marisa A. Wirth, Uta Passow, Jenny Jeschek, Ines Hand, and Detlef E. Schulz-Bull. Partitioning of oil compounds into marine oil snow: Insights into prevailing mechanisms and dispersant effects. Marine Chemistry, 206:62–73, 2018. doi: 10.1016/j.marchem.2018.09.007.

Tingting Yang, Kelly Speare, Luke McKay, Barbara J. MacGregor, Samantha B. Joye, and Andreas Teske. Distinct Bacterial Communities in Surficial Seafloor Sediments Following the 2010 Deepwater Horizon Blowout. Frontiers in Microbiology, 7, 2016. doi: 10.3389/fmicb.2016.01384.

Mark B. Yunker, Robie W. Macdonald, Peter S. Ross, Sophia C. Johannessen, and Neil Dangerfield. Alkane and PAH provenance and potential bioavailability in coastal marine sediments subject to a gradient of anthropogenic sources in British Columbia, Canada. Organic Geochemistry, 89–90:80–116, 2015. doi: 10.1016/j.orggeochem.2015.10.002.

Lei Zhang, Yang Cao, Nicholas S. Colella, Yong Liang, Jean-Luc Brédas, K. N. Houk, and Alejandro L. Briseno. Unconventional, Chemically Stable, and Soluble Two-Dimensional Angular Polycyclic Aromatic Hydrocarbons: From Molecular Design to Device Applications. Accounts of Chemical Research, 48(3): 500–509, 2015. doi: 10.1021/ar500278w.

Xiao Zhao, Wen Liu, Jie Fu, Zhengqing Cai, S. E. O’Reilly, and Dongye Zhao. Dispersion, sorption and photodegradation of petroleum hydrocarbons in dispersant-seawater-sediment systems. Marine Pollution Bulletin, 109(1):526–538, 2016. doi: 10.1016/j.marpolbul.2016.04.064.

